# EEG-GAN: A Generative EEG Augmentation Toolkit for Enhancing Neural Classification

**DOI:** 10.1101/2025.06.23.661164

**Authors:** Chad Williams, Daniel Weinhardt, Joshua Hewson, Martyna Beata Płomecka, Nicolas Langer, Sebastian Musslick

## Abstract

Electroencephalography (EEG) is a widely applied method for decoding neural activity, offering insights into cognitive function and driving advancements in neurotechnology. However, decoding EEG data remains challenging, as classification algorithms typically require large datasets that are expensive and time-consuming to collect. Recent advances in generative artificial intelligence have enabled the creation of realistic synthetic EEG data, yet no method has consistently demonstrated that such synthetic data can lead to improvements in EEG decodability across diverse datasets. Here, we introduce EEG-GAN, an open-source generative adversarial network (GAN) designed to augment EEG data. In the most comprehensive evaluation study to date, we assessed its capacity to generate realistic EEG samples and enhance classification performance across four datasets, five classifiers, and seven sample sizes, while benchmarking it against six established augmentation techniques. We found that EEG-GAN, when trained to generate raw single-trial EEG signals, produced signals that reproduce grand-averaged waveforms and time-frequency patterns of the original data. Furthermore, training classifiers on additional synthetic data improved their ability to decode held-out empirical data. EEG-GAN achieved up to a 16% improvement in decoding accuracy, with enhancements consistent across datasets but varying among classifiers. Data augmentations were particularly effective for smaller sample sizes (30 and below), significantly improving 70% of these classification analyses and only significantly impairing 4% of analyses. Moreover, EEG-GAN significantly outperformed all benchmark techniques in 69% of the comparisons across datasets, classifiers, and sample sizes and was only significantly outperformed in 3% of comparisons. These findings establish EEG-GAN as a robust toolkit for generating realistic EEG data, which can effectively reduce the costs associated with real-world EEG data collection for neural decoding tasks.

## Main

The complex functioning of the mind has long captivated humanity, and electroencephalography (EEG) has been one of the foremost tools for its investigation. EEG has provided insights into various aspects of cognitive functioning, including learning, perception, and attention [1, 2]. The decoding of EEG data has played a significant role in brain-computer interfacing [3], educational training [4], and neurofeedback conditioning [5]. Moreover, EEG data decoding has contributed to the assessment of brain health, leading to significant advancements in the tracking and diagnosis of psychiatric and neurodegenerative disorders [6, 7].

The key challenge in decoding EEG data lies in its low signal-to-noise ratio. This is primarily because the brain’s electrical activity must travel to the scalp, which significantly weakens the signal [6, 1]. The resulting signal is easily obscured by noise, including physiological artifacts, external electrical signals, and environmental electromagnetic fields. This issue can be addressed by training classification algorithms with large amounts of data, which allows a consistent neural signal to emerge from the noise [8]. However, collecting such large datasets is a time-consuming and expensive process, making classification with EEG often impractical [9]. Recent advancements in generative artificial intelligence (AI) offer a promising solution to this problem by generating synthetic EEG samples to augment the dataset. These samples can then be used to augment the training datasets required by data-intensive classifiers.

Generative AI can generate synthetic data samples that mimic the original data, thus enlarging the datasets available for classification algorithms. Critically, such methods can model the probability distribution of the original data, thus accounting for both signal and noise components. Standard generative AI models include generative adversarial networks (GANs [10, 11]), variational autoencoders (VAEs [12, 13]), and denoising diffusion probabilistic models (DDPMs [14, 15]). While these models have primarily been developed and applied for image generation, recent advancements suggest their potential for generating neural data, including EEG signals. Indeed, GANs [16, 17, 18, 19, 20, 21, 22, 23, 24, 25, 26, 27, 28, 29, 30, 31, 32], VAEs [33, 34, 35, 36, 37, 38, 39, 40, 41], and DDPMs [42, 43, 44, 45] have demonstrated the capacity to generate realistic EEG data and improve classification performance in neural decoding applications.

While the aforementioned studies highlight the potential of generative AI in EEG-based research, they have yet to demonstrate a clear superiority over standard EEG data augmentation techniques, which limits their broad adoption within the neuroscience community. A critical limitation of prior applications of generative AI for EEG data augmentation is that they were often confined to a single dataset, classification algorithm, EEG system, and EEG processing pipeline, raising reservations about their generalizability across experimental settings. Furthermore, these studies often lack rigorous comparisons with benchmark augmentation techniques—such as transformation-based methods commonly used in neuroscience [46] and other prominent generative models, such as VAEs [47]. Another overlooked factor is the impact of data augmentation across varying sample sizes, which are highly variable in real-world applications. For instance, one of the most comprehensive evaluations to date, conducted by Fahimi et al. [20], demonstrated improved classification performance using a deep convolutional GAN on two datasets and two classifiers, compared against three benchmark techniques. However, their analysis was limited to a single EEG system, EEG processing pipeline, and sample size, leaving the broader applicability of their findings unclear. Finally, and perhaps most importantly, existing approaches often lack open accessibility, user-friendly interfaces, and adequate documentation, posing significant barriers to their widespread adoption within the neuroscience community. In the current study, we introduce EEG-GAN—an open-source toolkit featuring a generative adversarial network (GAN) for EEG data augmentation (see EEG-GAN’s Documentation [48] for user-friendly tutorials accompanied with detailed documentation). We evaluate the capability of this toolkit for augmenting EEG data and enhancing classification performance across an extensive range of contexts. Specifically, we assessed GAN-augmentation effects across four datasets, five classifiers, and seven sample sizes and compared our findings to six benchmark augmentation techniques. The four datasets represented distinct cognitive domains—learning, cognitive control, perception, and attention—collected independently by three research groups in Canada (learning [49]), Switzerland (cognitive control [50]), and the United States (perception and attention [51]), see Figure 1. These groups recorded their datasets using different EEG systems—ActiCHamp (Brain Products), Geodesic Hydrocel Sensor Net (Electrical Geodesics), and ActiveTwo (BioSemi), respectively—and processing pipelines. The datasets elicited key ERP components, including the reward positivity, N2-P300 complex, N170, and N2PC, alongside their corresponding frequency components. The five classifiers range from deep-learning models to tree-based models, each performing binary classifications across all datasets. We also probe seven sample sizes commonly found in EEG research. Specifically, we assess a range of realistic sample sizes for EEG research to ensure the practical application of EEG-GAN—specifically, sample sizes of 5, 10, 15, 20, 30, 60, and 100 participants. Whereas smaller sample sizes, ranging from 5 to 20 participants, are common when investigating specialized populations (e.g., clinical populations, early childhood), moderate sample sizes, ranging from 30 to 100 participants, are more common when investigating the general population. Thus, a broad range of sample sizes allows us to examine the applicability of EEG-GAN across varying data constraints. Finally, we compared the performance of our GAN to six benchmarks, including five state-of-the-art transformation augmentations and one generative AI augmentation. Specifically, we included the transformation augmentations as they are standard practice in EEG research, and included a VAE as it is a leading method of generative augmentation in the field of EEG.

**Fig. 1.**
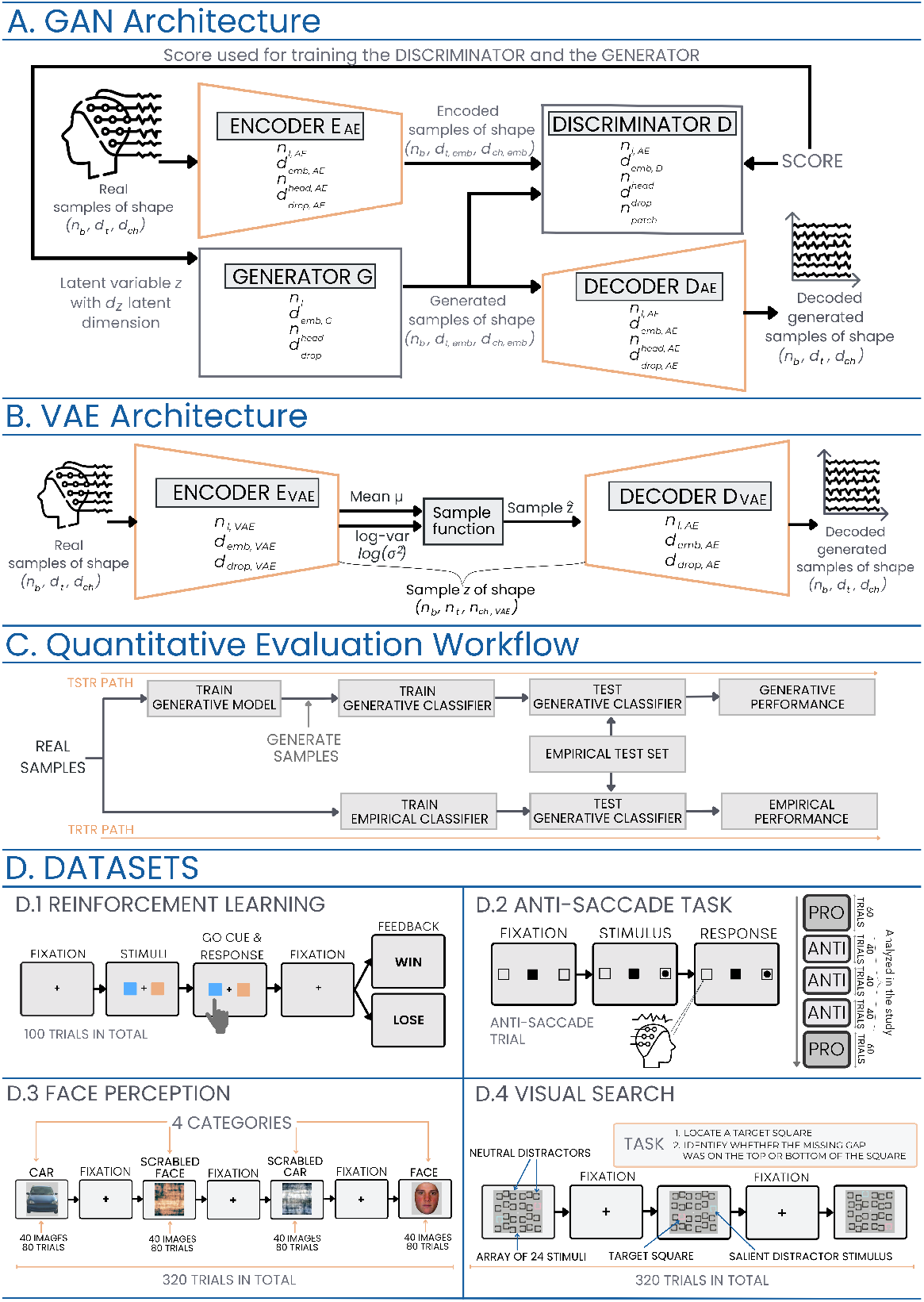
Overview of generative models and cognitive task datasets. A) Generative Adversarial Network (GAN) architecture, illustrating the interplay between encoder, generator, discriminator, and decoder components (see Generative Adversarial Network: EEG-GAN). B) Variational Autoencoder (VAE) structure, showing the encoding, sampling, and decoding processes (see Variational Autoencoder). C) Workflow diagram depicting the training and evaluation pipeline for generative models. D) Experimental datasets and paradigms: D.1) Reinforcement learning task, D.2) Antisaccade task, D.3) Face perception task, and D.4) Visual search task.

We found that EEG-GAN successfully generated realistic EEG samples that preserved cognitive phenomena. Using these samples for data augmentation improved classification performance across all datasets, classifiers, and sample sizes, with performance gains reaching up to 16%. The strongest effects were observed in smaller datasets of 30 participants or fewer—70% of which showed significant improvements. Furthermore, EEG-GAN outperformed all six benchmark augmentation techniques in 69% of comparisons.

## Results

### Quality of Generated EEG Data

To evaluate the quality of generated EEG samples produced by EEG-GAN, we employed two complementary approaches: (1) a visual and statistical comparison of synthetic versus empirical data across single-trial and aggregated metrics, and (2) a decoding-based evaluation that quantifies how well classifiers trained on synthetic data generalize to empirical data.

### Consistency with Empirical Data

Evaluations of generative AI models ensure that they can produce realistic EEG samples that reflect underlying cognitive phenomena across experimental settings. This includes both realistic single-trial samples, to which the models are trained to generate, and aggregated metrics (e.g., event-related potentials [ERPs], Fourier transforms [FFTs], time-frequency spectra [TFs]), which emerge through EEG processing pipelines. We first evaluated EEG-GAN’s ability to generate realistic EEG data and concomitant aggregated patterns across four datasets. We also compared samples generated by the EEG-GAN toolkit to those produced by an alternative generative AI (VAE) model (see Figure 1 and Methods).

The toolkit produced single-trial samples that resembled original samples from the datasets, whereas the VAE produced smoother samples that were more visually dissimilar to the raw data (see Figure S1). Processed samples from both models produced grand-averaged ERP waveforms that strongly correlated with empirical grand-averaged ERP waveforms (see Figure 2 and Table 1). Moreover, grand-averaging synthetic EEG data of both models produced the signature ERP components—the reward positivity, N2-P300 complex, N170, and N2PC—resembling those of the original datasets (see Figure 2), even though the models were not explicitly trained to generate grand-averaged patterns. Lastly, the single-trial samples produced similar grand-averaged time-frequency and frequency distributions for all datasets that exhibited typical features found in frequency decompositions of EEG data, such as the 1*/f* slope (see Figure 3 and Supplemental Figures S2, S3).

**Table 1.** Pearson correlations between the grand averaged difference waveforms of the empirical EEG data and the synthetic data generated by the GAN and VAE models. The difference waveforms were smoothed using a moving average filter with a window size of 5% of the total number of samples.

| Dataset | GAN | VAE |
| --- | --- | --- |
| **Reinforcement Learning (E1)** | 0.92 | 0.97 |
| **Reinforcement Learning (E8)** | 0.97 | 0.96 |
| **Antisaccade** | 0.55 | 0.87 |
| **Face Processing** | 0.86 | 0.09 |
| **Visual Search** | 0.49 | 0.44 |
| **Average** | **0.76** | **0.67** |

**Fig. 2.**
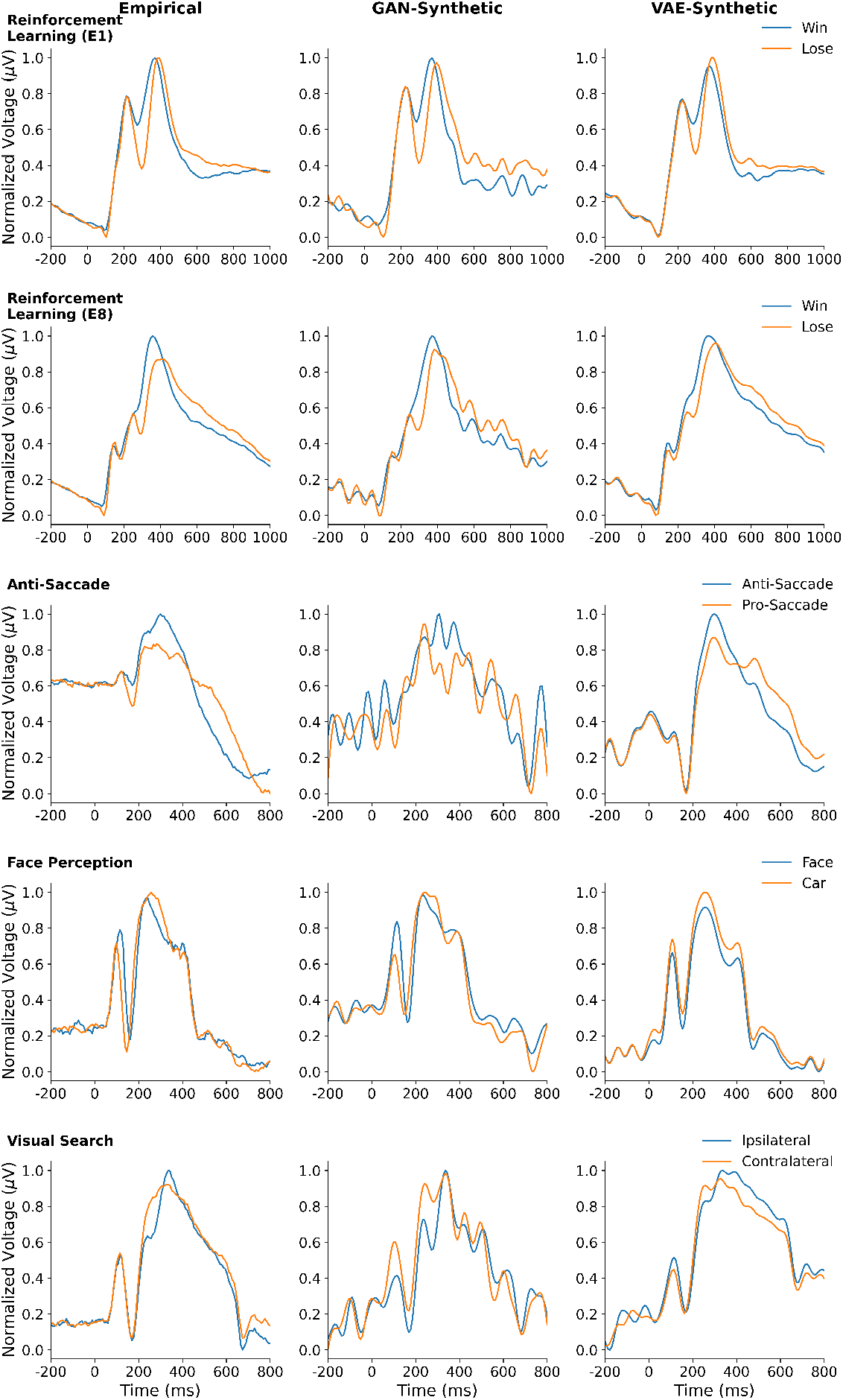
Grand-averaged ERP waveforms of processed empirical or synthetic single-trial data generated by the GAN and VAE models. Reinforcement Learning (E8) displays data at electrode POz, and the other datasets display electrodes as described in the methods.

**Fig. 3.**
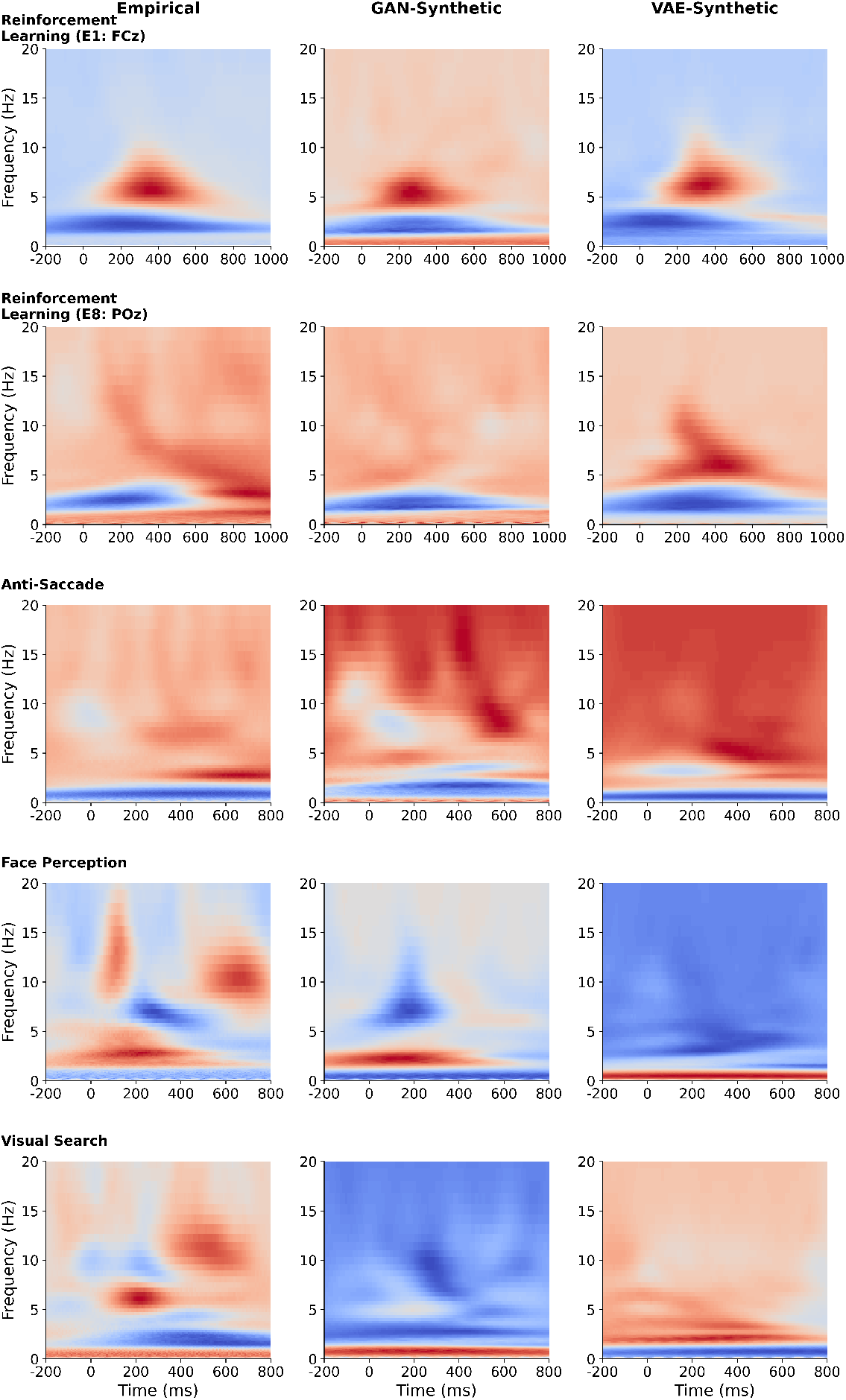
Gand-averaged time-frequency data from empirical and synthetic samples produced by GAN and VAE models.

### Evaluation of Decoding Performance

The efficacy of the EEG-GAN toolkit was further assessed with decoding analyses comparing the *Train on Synthetic, Test on Real* (TSTR) with *Train on Real, Test on Real* (TRTR) performances (see Figure 1C). TSTR assesses how well generated data generalizes to empirical data by training a classifier with generated samples and evaluating it on held-out empirical samples. Comparing TSTR performance with TRTR performance provides a direct evaluation of the quality of the generated data, where similar performance indicates that the generated data effectively captures the underlying distribution of the empirical data. Training the classifier just with GAN-generated data achieved a two-way classification performance comparable to when it was trained just with empirical data (e.g., 66% versus 65%, respectively, see Table 2). In contrast, training the classifier with data generated from an alternative (VAE) model resulted in diminished performance compared to when trained with empirical data (e.g., 57% versus 65%, respectively, see Table 2).

**Table 2.** Two-way classifier accuracy in % (mean [SD]) comparison between empirical (EMP) data (TRTR analysis), GAN-generated data (TSTR analysis), and VAE-generated data (TSTR analysis) for each dataset. Results illustrate the efficacy of GAN in achieving comparable two-way performance to empirical data, while VAE-generated data shows reduced classifier performance.

| Dataset | EMP | GAN | VAE |
| --- | --- | --- | --- |
| **Reinforcement Learning (E1)** | 71 (2) | 68 (1) | 65 (2) |
| **Reinforcement Learning (E8)** | 70 (2) | 69 (1) | 62 (3) |
| **Anti-Saccade** | 69 (4) | 60 (1) | 49 (4) |
| **Face Perception** | 65 (16) | 72 (3) | 59 (5) |
| **Visual Search** | 52 (7) | 60 (2) | 50 (8) |
| **Average** | **65 (10)** | **66 (5)** | **57 (8)** |

### Utility of Generated Samples for Data Augmentation

We next evaluated the utility of the generated samples for data augmentation, assessing whether incorporating synthetic data into classifier training could enhance decoding performance on empirical held-out datasets. Augmenting EEG data using GANs significantly enhanced classification performance across all four datasets, five classifiers, and the majority of sample sizes (see Figure 4 and Supplemental Figures S4-S8).

**Fig. 4.**
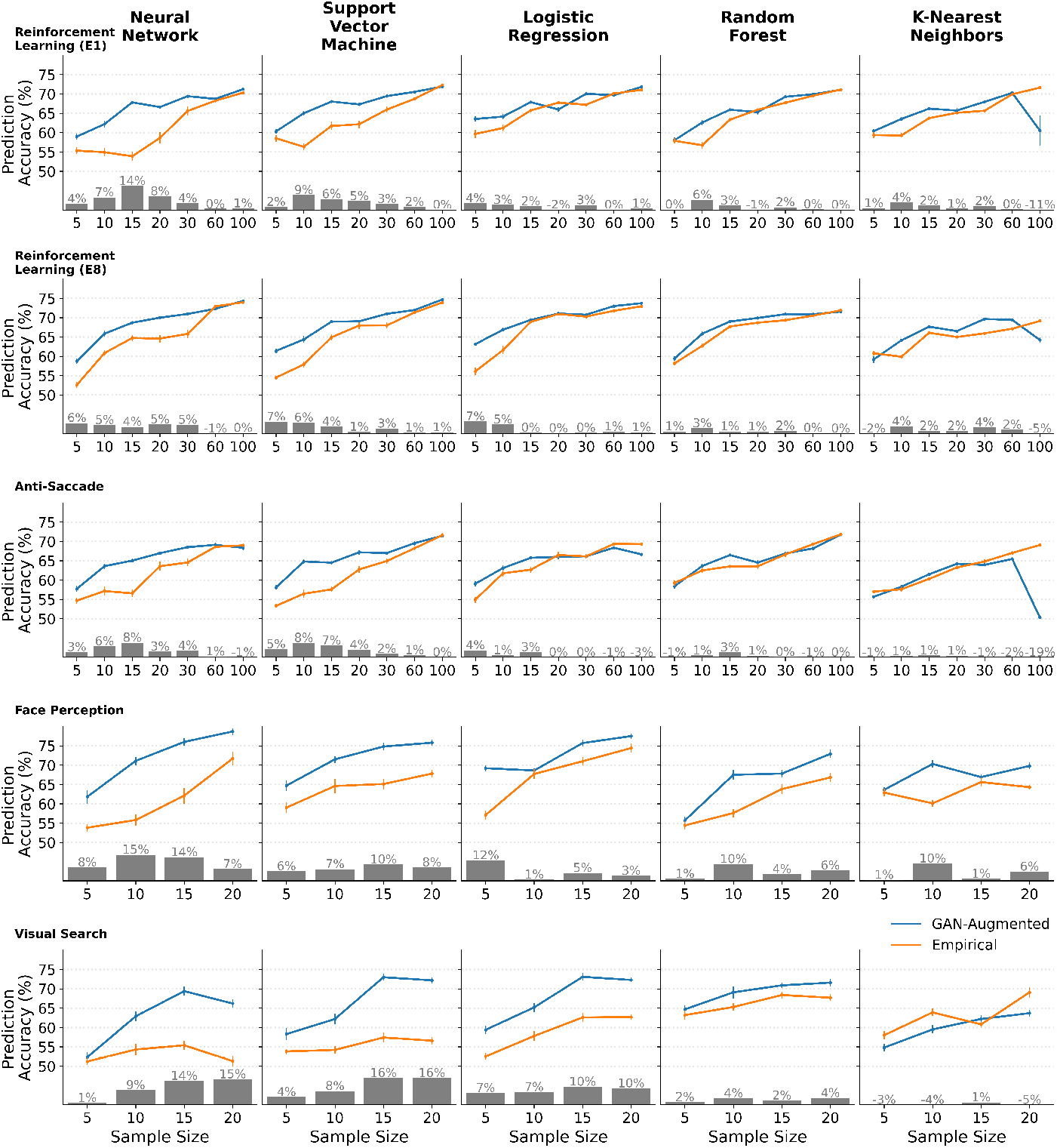
Impact of GAN data augmentation on EEG classification performance across four datasets, five classifiers, and four to seven sample sizes. Lines illustrate two-way classification performance for classifiers trained on empirical data alone (orange) and on additional synthetic data generated by EEG-GAN (blue). Data augmentation resulted in enhanced classification performance on empirical held-out data, with improvements of up to 16% two-way classification performance. The effects were consistent across datasets and varied among classifiers. GAN augmentations were most effective for smaller sample sizes (30 and below), where they enhanced 70% of classification analyses and only impaired 4% of analyses. Error bars represent standard error of the mean. The grey bar plots depict decoding performance improvements after including synthetic samples in the training set.

GAN augmentations significantly impacted two-way classification performance, resulting in an increase in prediction accuracy of up to 16%. Specifically, GAN augmentations consistently improved classification performance across four datasets and five classifiers, with greater benefits for certain classifiers (e.g., neural networks, logistic regression) and smaller sample sizes of 30 and below. Indeed, for sample sizes of 30 and below, GAN augmentations significantly enhanced 70% of classification analyses (60% across all sample sizes) and only significantly impaired 4% of analyses (8% across all sample sizes). For instance, when augmenting a dataset containing five participants, classifier performance was equivalent to that of an empirically trained model with four times the number of participants.

We also benchmarked EEG-GAN against five state-of-the-art transformation-based augmentation methods and one generative augmentation method. In the transformation-based methods, samples were directly altered—for example, the time reverse transformation inverted the temporal dimension of selected samples, making classifiers more resilient to effects that vary in time. For the generative method, we generated samples using a VAE and appended them to the training datasets, following the same procedure as with the GAN-generated samples.

EEG-GAN robustly outperformed all conventional augmentation benchmarks (see Supplemental Figures S9-S13). As this evaluation involved numerous comparisons, we also provide an interactive plot that allows readers to control which contrasts are displayed: interactive supplemental figure [48]. GAN augmentation elicited greater classification enhancements than the six benchmark augmentation techniques for 69% of analyses across sample sizes 30 and below (63% across all sample sizes) and was only outperformed by 3% of comparisons (5% across all sample sizes).

## Discussion

We present EEG-GAN, an open-source data augmentation toolkit that leverages generative adversarial neural networks (GANs) to enhance neural decoding performance. By generating single-trial samples, EEG-GAN is adaptable to a wide range of experimental paradigms, making it a versatile resource for EEG researchers and practitioners. The toolkit is available as an easy-to-use Python package, developed and maintained according to professional software engineering standards (Documentation [48]). We conducted a comprehensive evaluation, confirming the utility of EEG-GAN across four datasets with distinct EEG systems and preprocessing pipelines, five classification algorithms, and seven training set sizes.

Our evaluation revealed that EEG-GAN generates realistic single-trial samples, which, when processed through standard EEG analysis pipelines, reproduce key grand-averaged patterns, such as ERPs and time-frequency spectra, across all datasets. Notably, the model was not explicitly trained to generate these aggregate features, yet the synthetic data reliably captured characteristic patterns that emerge at the grand-average level. These include hallmark ERP components such as the reward positivity, N2-P300 complex, N170, and N2PC—spanning diverse cognitive domains including reinforcement learning, cognitive control, perception, and attention. This suggests that EEG-GAN captures distributional features of the data along with key qualitative patterns that emerge from them.

Moreover, the generated samples demonstrated practical utility by significantly enhancing classifier performance on empirical held-out datasets when included in the training data. This improvement was achieved without the addition of new real data, indicating that the synthetic samples act as plausible interpolations within the original data distribution. By enriching the data distribution, the generated samples help constrain the hypothesis space of the classifier, enhancing its generalization performance.

The demonstrated ability of EEG-GAN to generate realistic and high-quality EEG data opens new avenues for studying cognitive processes while reducing the need for extensive and costly data collection [2]. By enabling researchers to achieve decoding performance comparable to that obtained from dozens of additional participants, EEG-GAN offers a cost-effective and computationally efficient alternative (see Figure 4). This capability is particularly valuable in studies and contexts where only limited EEG data can be collected, such as in clinical settings where participant availability or practical constraints often limit data collection. In such scenarios, EEG-GAN’s data augmentation could improve the robustness of neural decoding analyses. Additionally, the enhanced classification performance enabled by GAN-augmented data has the potential to support more accurate and reliable decoding applications across domains of study [3, 4, 5].

While our study provides valuable insights, it also reveals promising directions for future research. Expanding comparisons beyond the VAE benchmark included in this study to other generative augmentation methods, such as GAN and DDPM architectures, could provide a more comprehensive understanding of their relative strengths, particularly using recent generative models [42]. Additionally, while our study demonstrates the potential of GAN-augmented EEG data in enhancing classification performance, further research is needed to explore its practical applications in real-world settings and with mobile EEG technology [6, 52]. Such efforts will not only refine our understanding of generative augmentation techniques but also pave the way for their integration into real-world EEG applications.

In conclusion, we introduced the EEG-GAN toolkit for augmenting EEG data and examined its utility for EEG decoding. Our findings demonstrate that EEG-GAN can generate realistic single-trial samples that accurately reproduce grand-averaged patterns, such as ERPs and time-frequency spectra, and significantly enhance classification performance in neural decoding tasks. By providing a cost-effective solution for data augmentation, EEG-GAN addresses critical challenges in EEG research, particularly in scenarios where large-scale data collection is impractical or infeasible, such as mobile or clinical EEG applications. Beyond demonstrating its utility across various datasets, the evaluation of EEG-GAN lays a foundation for future explorations into generative augmentation methods, highlighting their potential applications in neuroscience and neurotechnology.

## Methods

### Data and Script Availability

All data, simulations, and analysis scripts used here are available at https://autoresearch.github.io/EEG-GAN [1].

### Datasets

#### Dataset 1: Reinforcement Learning

In line with Williams, Weinhardt, and colleagues [2], we evaluated EEG-GAN on a large open-source dataset provided by Williams and colleagues [3] that contains 500 participants performing a two-armed bandit task. For evaluation analyses (see Quality of Generated EEG Data), all participants were included, and for classification analyses (see Utility of Generated Samples for Data Augmentation), participants were split into 100 training, 200 validation, and 200 test sets (see Supplemental Table 5). In this study, participants completed a simple two-armed bandit reinforcement learning task while EEG data were recorded using a standard 32- or 64-channel system. We will here provide a brief description of the study, including the task, EEG setup, and EEG processing, but see Williams et al. [3] for a detailed description.

The two-armed bandit task is a canonical paradigm for assessing the neural correlates of reinforcement learning. This task had participants decide which of two stimuli was more often rewarding through trial and error (see Figure 1D1). On each trial, participants saw two coloured squares and had to decide between them, receiving binary win or lose feedback. For each pair of coloured squares, one of the squares had a 60% chance of providing win feedback, and the other had a 10% chance of providing win feedback. Participants saw each pair of coloured squares twenty times. They saw five pairs of stimuli, resulting in 100 trials per participant, with roughly equal counts of the win and lose conditions (47% of trials were within the win condition). The task elicits three neural components of reward processing: the reward positivity event-related potential, delta oscillations, and theta oscillations [3]. Here, the goal was to classify between win and lose conditions.

Participant EEG data were recorded at a 500 Hz sampling rate using a standard 32 or 64 electrode EEG system (ActiCAP, Brain Products, GmbH, Munich, Germany). Williams and colleagues [3] provide all pre-processed data, which are used here. See EEG Pre-Processing for details on how they processed the data. These processing steps resulted in each sample containing 600 data points. For the current study, we downsampled these data to 100 data points (an effective sampling rate of 83.33 Hz) and normalized them within each trial to a range of 0 to 1. For this dataset, we conducted all analyses twice: once using a single electrode, FCz, and again using eight electrodes with full coverage across the scalp: FCz, F3, F4, C3, C4, P3, P4, and POz. For the latter, this entailed training generative models using data from all eight electrodes and subsequently generating data specific to each electrode.

#### Dataset 2: Anti-Saccade

As a second dataset, we used the large open-source EEGEyeNet dataset provided by Kastrati et al. [4] with 329 recordings of participants conducting a series of gaze tasks. Here, we focused on the anti-saccade task, in which participants looked left or right in accordance with or in opposition to a spatial cue, while EEG and eye-tracking data were recorded. Initially, 129 participants underwent two experimental sessions following the same EEG protocol to evaluate the test-retest reliability of all dependent measures. However, we selected only the data from each participant’s first session for the current analyses. Consequently, this approach provided simultaneous EEG and eye-tracking recordings from 200 participants. The data of one of these participants was missing due to technical issues, and we excluded ten other participants with excessively noisy grand-averaged ERPs post-processing (see [4]). For evaluation analyses (see Quality of Generated EEG Data), we used the 189 participants who were deemed acceptable, whereas for classification analyses (see Utility of Generated Samples for Data Augmentation), we used all 199 participants. For classifications, data were randomly split into training, validation, and test sets of 100, 49, and 50 participants, respectively (see Supplemental Table 5). A brief description of the task, EEG setup, and EEG processing are described below, but see Kastrati et al. [4] for detailed explanations.

The anti-saccade task requires participants to look left or right in response to a spatial cue (see Figure 1D2). On each trial, a cue (i.e., a dot) appeared on the left or right side of the visual field, and participants were either to move their gaze towards (congruent) or away from (incongruent) the cue. The participants engaged in 120 trials for each of the congruent and incongruent conditions, and the direction of their gaze (left, right) was equiprobable. We only used trials where the participants looked left, leaving us with 60 trials per participant. Of these, only trials where the participants performed correctly were used. The anti-saccade task elicits an N2-P300 ERP complex that reflects conflict monitoring and response inhibition, respectively [5]. The goal here was to classify between congruent and incongruent conditions.

Participant EEG data were recorded at 500 Hz using a standard 128 Geodesic Hydrocel EEG system (Electrical Geodesics Inc., Oregon, USA). We used the raw data provided by Kastrati et al. [4], see EEG Pre-Processing for details on how the data were processed. These steps resulted in data samples with 500 data points, which we then downsampled to 125 data points (i.e., 125 Hz), and then we normalized each trial to be between 0 and 1. We extracted an averaged central electrode, including Cz, E6, E7, E13, E106, and E112.

#### Dataset 3: Face Perception

The third dataset was drawn from ERP CORE, a gold-standard database of ERP experiments [6]. ERP CORE contains EEG data of 40 participants who conducted six tasks, eliciting seven distinct ERP components. The task investigated in this dataset was the face perception task, wherein participants viewed images of faces and cars, as well as their scrambled variants, while EEG data were recorded on a standard 30-electrode system. The authors of ERP CORE removed three of the 40 participants from their analyses due to high artifact rejection rates. For evaluation analyses (see Quality of Generated EEG Data), we included the remaining 37 participants with acceptable data, but for classification analyses (see Utility of Generated Samples for Data Augmentation), all 40 participants were used and were randomly split into training, validation, and test sets containing 20, 10, and 10 participants, respectively (see Supplemental Table 5). We will again provide brief descriptions of the task, EEG setup, and EEG processing, but see Kappenman et al., [6] for detailed descriptions of each. The face perception task involved participants viewing images of faces and cars, along with their scrambled counterparts (see Figure 1D3). The participants were tasked with indicating, on each trial, whether the image contained an object (face or car) or a texture (scrambled object). Each category (face, car, scrambled face, scrambled car) contained 40 images, and each image was presented twice throughout the experiment, resulting in a total of 320 trials. The face perception task elicits an N170 ERP component that exhibits differently for faces than non-faces. As such, our study was interested in classifying faces versus cars (i.e., non-faces) and thus did not include trials of scrambled stimuli. The goal here was to classify between face and non-face conditions.

EEG data were recorded at 1024 Hz using a standard 30-electrode BioSemi system (Biosemi B.V., Amsterdam, the Netherlands). Kappenman et al. [6] provided all data in a fully pre-processed state, which was used here; see EEG Pre-Processing for details on how the data were processed. These processing steps resulted in each sample containing 256 datapoints, from which we then downsampled the data to 128 datapoints (i.e., 128 Hz) and normalized the result to a range of 0 to 1 for each trial. We also reduced the data to the electrode of interest, signified in ERP CORE for this task, namely PO8.

#### Dataset 4: Visual Search

The fourth dataset was also drawn from ERP CORE [6]. This dataset contains EEG data from 40 participants conducting a visual search task, where participants were asked to locate an oddball target stimulus among an array of distractor stimuli. The authors of ERP CORE removed five of the 40 participants from their analyses as they had high artifact rejection rates. As such, evaluation analyses (see Quality of Generated EEG Data) were conducted on the 35 acceptable participants, but classification analyses (see Utility of Generated Samples for Data Augmentation) used all 40 participants, which were randomly split into a training dataset of 20 participants, a validation dataset of 10 participants, and a test dataset of 10 participants (see Supplemental Table 5). We provide brief descriptions of the task, EEG setup, and EEG processing, but full details can be found in Kappenman et al. [6].

The visual search task has participants searching through an array of 24 stimuli (see Figure 1D4). The stimuli used here were the outline of squares with a missing gap on one side. The goal of the task was for participants to locate a target square, which was either coloured pink or blue, and identify whether the missing gap was on the top or bottom of the square. One of the other squares was a salient distractor stimulus, which was coloured in pink or blue opposite the target and also had a gap on the top or bottom. The remaining 22 neutral distractor stimuli were coloured in black and had their gaps either on the left or right side of the square. The participants conducted this task for 320 trials. The visual search task elicits an N2pc (N200-posterior-contralateral) ERP component that reflects directed attention. This component is larger on the contralateral side of recording, depending on the target location (e.g., if the target is in the left visual hemisphere, there is an enhanced neural response on the right hemisphere of the brain). Conditions in this task occur within a trial by comparing neural activity in the contralateral hemisphere to the ipsilateral hemisphere. Here, we sought to classify contralateral from ipsilateral activity. Here, the goal was to classify between contralateral and ipsilateral conditions.

EEG data were recorded and processed in the same manner as dataset 2 (see section Dataset 2: Anti-Saccade and EEG Pre-Processing) except that the data were averaged to joint mastoid electrodes and the electrodes of interest were PO7 and PO8 (which would each reflect either ipsilateral or contralateral conditions dependent on the target location on a given trial).

### Augmentation Techniques

Next, we will outline the GAN architecture used in the evaluation study. See supplemental materials for descriptions of the Benchmark Augmentation Techniques.

### Generative Adversarial Network: EEG-GAN

Preliminary work suggests that augmenting classifier training sets with synthetic EEG samples produced by generative adversarial networks (GANs) can improve classification performance [2]. In this study, we present a novel GAN architecture, EEG-GAN, and demonstrate its efficacy in EEG data augmentation across an extensive range of experimental contexts.

The GAN [7] consists of two opposing networks—the generator *G* and the discriminator *D*. Whereas the generator produces samples, the discriminator distinguishes between real and generated samples. During training, the GAN is deemed successful in generating realistic samples if the discriminator can not differentiate between real and generated samples.

Specifically, the generator *G* produces a sample *ŷ* from a vector *z*, which is randomly sampled according to a probability distribution *P*_*z*_ as determined by the generator parameters *θ*_*G*_:

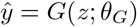

The discriminator *D* receives either real samples *y* or generated samples from the generator *ŷ* and assigns a validity score *v*, which amounts to a probability that the sample was drawn from the empirical dataset relative to the discriminator parameters *θ*_*D*_:

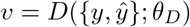

The GAN developed by Goodfellow and colleagues [7] encounters multiple obstacles, including training instability and mode collapse, which limit its application.

However, it provides a framework for improvement. To address these issues, we modified the traditional GAN architecture in several ways. First, we replaced the traditional GAN’s objective function with a Wasserstein distance [8]:

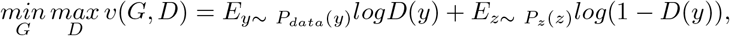

where the discriminator’s function maximizes the distance:

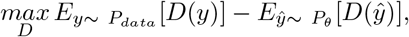

and the generator’s function minimizes the distance:

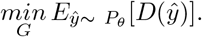

We further included a gradient penalty (GP) for each update of the discriminator’s parameters *θ*_*D*_, thus penalizing high gradients [9]. Next, due to the temporal dependencies of EEG data, we implemented both the generator and discriminator as transformer models [10, 11]. Finally, we applied a conditional GAN that incorporated conditional labels in both the generator and discriminator, allowing the GAN to contextualize the data based on pre-specified experimental conditions [12].

While these modifications suggested that the architecture can generate realistic EEG samples [2], we theorized that the noisy nature of EEG data is decreasing training stability, resulting in long training times and unstable results. As such, we have further improved the GAN architecture by embedding an autoencoder (AE). That is, instead of feeding the time-series data directly to the GAN, we provide it with an encoding of the EEG data produced by the AE. This structure results in quicker and more stable training as it decouples feature extraction from generation, simplifying the training process.

The AE is composed of two networks, an encoder *E*_*AE*_ and a decoder *D*_*AE*_—each being a transformer. Data passing through the AE is encoded into an embedded form *y*_*emb*_ with *n* units relative to the AE encoder parameters 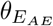:

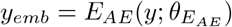

The AE embedding is then reconstructed back into the original data space *ŷ*_*recon*_ by the decoder as determined by the AE decoder parameters 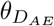:

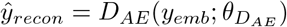

The autoencoder is trained to minimize the mean squared error between the input data *y* and the reconstructed data *ŷ*_*recon*_:

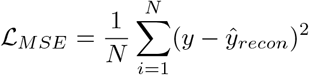

We trained the AE with an ADAM optimizer with a *l*_*r*_ = 0.0001 and decay rates of *β*_1_ = 0, *β*_2_ = 0.9.

The AE is embedded into the GAN architecture by having the generator produce data akin to the AE’s embedded layer (*y*_*emb*_) rather than the original time-series (*y*):

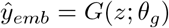

Likewise, the discriminator receives either real samples that are encoded into the embedded layer *y*_*emb*_ or fake samples that are generated to reflect the embedded layer *ŷ*_*emb*_:

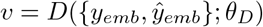

Once training is completed, EEG samples are generated by first having the generator produce embedded layer data *ŷ*_*emb*_ and then having the autoencoder decode this data into the original EEG data space:

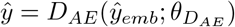

GAN training is performed on trial-level EEG data with an ADAM optimizer with *l*_*r*_ = 0.0001 and decay rates of *β*_1_ = 0, *β*_2_ = 0.9 (Gulrajani et al., [9]). The generator-to-discriminator training frequency is set to 1 : 5 (Arjovsky [8]).

## Analyses

### Quality of Generated EEG Data

We evaluated both qualitatively and quantitatively whether EEG-GAN, as well as a benchmark generative model (VAE), can accurately generate EEG data across different experimental paradigms and cognitive phenomena.

#### Consistency with Empirical data

For qualitative evaluations, we trained the generative models on all participants (Dataset 1: 500 participants; Dataset 2: 199 participants; Dataset 3: 37 participants; Dataset 4: 35 participants) for 2,000 epochs and produced 2,500 samples per condition (Dataset 1: win, lose; Dataset 2: anti-saccade, pro-saccade; Dataset 3: face, car; Dataset 4: contralateral, ipsilateral). We then visually compared these generated samples to empirical samples along four traditional representations of EEG data: 1) trial-level EEG data, 2) grand-averaged ERPs, 3) grand-averaged frequency transforms, and 4) grand-averaged time-frequency transformations. For example, in Dataset 1’s reinforcement learning task, successfully generated data would demonstrate realistic trial-level EEG data, the reward positivity ERP component, and delta and theta effects in the frequency/time-frequency dimension [3]. We also conducted correlational analyses on the grand-averaged difference waveforms. Specifically, for each empirical dataset, we grand-averaged conditional waveforms and then created difference waveforms. We conducted the same procedure for each of the GAN-generated and VAE-generated samples. We then smoothed each of these grand-averaged difference waveforms with a moving average window spanning 5% of the total data length. We then correlated each of these generated difference data with the empirical difference data, allowing for a comparison of the empirical patterns with those generated by the GAN and VAE.

#### Evaluation of Decoding Performance

Additionally, we conducted quantitative evaluations by comparing the performance between Train on Synthetic, Test on Real (TSTR) analyses and Train on Real, Test on Real (TRTR) analyses [13]. The TSTR method ensures that synthetic data can serve as an effective substitute for empirical data by training a classifier on synthetic samples and evaluating its performance on held-out empirical samples. This approach tests the ability of synthetic data to generalize to empirical data, thereby reflecting how well it captures the underlying distribution of the empirical dataset. Equal performance of the TSTR and TRTR methods indicates that the synthetic data is as informative as the empirical data when predicting held-out empirical samples. To conduct these analyses, we first trained each generative model on the training datasets (Datasets 1 & 2: 100 participants, Datasets 3 & 4: 20 participants). We produced 2,500 samples of data for each of the corresponding conditions. These data were then averaged in chunks of 50, resulting in 50 synthetic participants, each with an averaged time series per condition. We then trained simple neural networks (see using the empirical training dataset, the GAN-generated training dataset, and the VAE-generated training dataset. The performance of the neural network classifiers was then determined by their ability to predict the labels of the corresponding dataset’s test set (Dataset 1: 200 participants, Dataset 2: 100 participants, Dataset 3: 10 participants, Dataset 4: 10 participants). The classifier training and testing procedure was conducted ten times for each dataset and data training type (empirical, GAN, VAE).

### Utility of Generated Samples for Data Augmentation

#### Experimental Factors

The primary focus of the current study was to assess the degree to which synthetic samples produced by EEG-GAN enhance EEG decoding performance when included in the training set of traditional classifiers. Furthermore, we sought to examine whether this decoding enhancement is robust across a large range of contexts, as determined by four experimental factors:

- **Datasets (4)**: Reinforcement Learning, Anti-Saccade, Face Perception, Visual Search. Combined, these datasets encompass 4 ERP components, 3 EEG Systems, and 3 distinct EEG processing pipelines.
- **Augmentation Techniques (7)**: EEG-GAN, VAE, Oversampling, Gaussian Noise, Time Reverse, Polarity Reverse, Data Smoothing
- **Sample Sizes (4 - 7)**: 5, 10, 15, 20, 30, 60, 100
- **Classifiers (5)**: Neural Networks, Support Vector Machines, Logistic Regression, Random Forest, K-Nearest Neighbours.

Our first factor is the **dataset** used for testing. This study utilized four EEG datasets from participants performing standard psychological tasks. Specifically, we examined the effects of EEG-GAN augmentation on classification performance in a reinforcement learning task, an anti-saccade task, a face perception task, and a visual search task (see Section Datasets). These are all open-source datasets from distinct laboratories using various EEG systems and EEG processing methodologies (see the dataset descriptions above).

Our second factor includes a range of **augmentation techniques**. Our goal in this study was to evaluate EEG-GAN, and we also aimed to compare its data augmentation capabilities with those of traditional methods. As such, we included one generative augmentation benchmark (see section Generative Adversarial Network: EEG-GAN)— a variational autoencoder—and five state-of-the-art transformation augmentation benchmarks (see section Transformation Augmentation Techniques)—oversampling, Gaussian noise, time reverse, polarity reverse, and data smoothing (see Transformation Augmentation Techniques).

Our third experimental factor is the **sample sizes** used to train and test the GANs. Although some of the datasets we used contain large amounts of data (e.g., Dataset 1 has 500 participants), we aimed to investigate augmentation performance across a broad range of typical sample sizes commonly found in EEG studies. As such, we randomly split our training sets into subsets of varying sample sizes for training. For datasets 1 & 2, our training dataset contained 100 participants, and we extracted subset sample sizes of 5, 10, 15, 20, 30, 60, and 100 participants. For datasets 3 & 4, our training datasets contained 20 participants, and we extracted subset sample sizes of 5, 10, 15, and 20 participants. We repeated this process five times for each sample size. As such, for datasets 1 & 2, we had 35 training datasets (five per seven sample sizes), and for datasets 3 & 4, we extracted 20 training datasets (five per four sample sizes). The randomization process was independent across all subsets, so participants would overlap across the training subsets. Furthermore, the subsets that were similar in size to the full training dataset (that is, 100 participants for datasets 1 & 2 and 20 participants for datasets 3 & 4) would contain all the participants included in training and thus would be identical. Nonetheless, we still extracted these datasets five times and trained five separate generative models.

Our fourth and final experimental factor was the **classifiers** for which we evaluated the effects of data augmentation. We assessed whether EEG-GAN augmentation enhanced classification performance on simple neural networks (NN), support vector machines (SVM), logistic regressions (LR), Random Forests (RF), and K-Nearest Neighbours (KNN). Each of these classifiers was implemented using Scikit-learn [14] and included a grid search of meta-parameters (see Supplemental Table 5). The input data for each classifier was participant-averaged full time-series data for the corresponding condition (reinforcement learning: win versus lose, anti-saccade: congruent versus incongruent, face perception: faces versus non-faces, visual search: ipsilateral versus contralateral, see Datasets). For example, the training dataset of 100 participants with two conditions in the reinforcement learning task would include 200 samples from each of the two generative models, EEG-GAN and VAE.

To assess the statistical significance of differences in classification performance between GAN-generated results and both empirical and benchmark performances, we applied a bootstrap resampling procedure within each dataset, classifier, and sample size. This procedure involved 10,000 iterations to generate null distributions of mean differences, centered at zero. These distributions were then used to compute p-values, representing the proportion of resampled mean differences that equaled or exceeded the observed mean difference of the corresponding comparison. Analyses were conducted using two-tailed tests with an *α* level of 0.05.

### EEG Data Generation and Augmentation

For the GAN and VAE, we trained each model, generated new samples, and used these during evaluation and classification analyses. Training a GAN first required training an autoencoder. The autoencoders were trained so that the encoder’s embedded layer roughly halved the size of the time-series dimension (Reinforcement Learning: 100 *⇒* 50, Anti-Saccade: 125 *⇒* 64, Face Perception: 128 *⇒* 64, Visual Search: 128 *⇒* 64). We also reduced this dimension to half the size of the reinforcement learning dataset with eight electrodes. These autoencoders were trained for 2,000 epochs and then embedded within the GAN, which was trained for 2,000 epochs. The VAE was also trained for 2,000 epochs.

For evaluation analyses, we generated a number of *synthetic participants* equal to the empirical datasets being compared (Dataset 1: 500 participants, Dataset 2: 199 participants, Dataset 3: 37 participants, Dataset 4: 35 participants). We generated 50 trials per condition (the minimum available trials across the datasets) for each synthetic participant and averaged these samples within condition and participant so that each synthetic participant had one participant-averaged sample per condition. For example, the reinforcement learning task had 500 participants within the evaluation analyses, and thus, the generative models each produced 500 synthetic participants by creating 50 trial-level samples per 2 conditions. Therefore, in this example, we generated 500 participants x 2 conditions x 50 trials = 50,000 trial-level samples.

We used each model to generate 5,000 samples per condition (e.g., win, lose in the reinforcement learning dataset), averaged in segments of 50 trials to create data of 100 *synthetic participants*. The number of synthetic participants used to create each augmented dataset for classifier training equaled the current sample size of interest, so that augmented datasets contained a 1:1 ratio between empirical and synthetic participants. For example, classifiers addressing a sample size of 5 would be trained on data from 5 empirical and five synthetic participants. Likewise, augmented datasets for a sample size of 100 participants would include 100 empirical and 100 synthetic participants. These synthetic participants were used as additional training samples during classification, and their impact was evaluated by comparing augmented classification performance to empirical (non-augmented) performance.

## Acknowledgements

The EEG-GAN framework is developed and maintained by members of the Autonomous Empirical Research Group. Chad Williams, Joshua Hewson, and Sebastian Musslick were supported by the Carney Brainstorm program at Brown University. Sebastian Musslick was also supported by Schmidt Science Fellows, in partnership with the Rhodes Trust. Nicolas Langer was supported by the Swiss National Science Foundation SNSF Projects (10001C_197480; 10001C_220048).

## Author Contributions

Chad Williams conceptualized the experiment, contributed to developing the EEG-GAN package, including its code and documentation, conducted the experiments, and authored the manuscript. Daniel Weinhardt also conceptualized the experiment, contributed to developing the EEG-GAN package, including its code and documentation, and reviewed the manuscript. Joshua Hewson contributed to the development of the EEG-GAN package, including its code and documentation. Martyna Beata Plomecka collected and processed data and reviewed the manuscript. Nicolas Langer reviewed the manuscript. Sebastian Musslick conceptualized the experiment, supervised the project, and reviewed the manuscript.

## Supplementary information

### EEG Pre-Processing

#### Reinforcement Learning

1. Data with a 64 electrode system were reduced to 32 electrodes.
2. Noisy or damaged electrodes were removed.
3. Data were re-referenced to an average mastoid reference.
4. Data were filtered using a 0.1 to 30 Hz Butterworth filter (order 4).
5. Data were filtered using a 60 Hz notch filter.
6. ICA was employed to identify and correct eye blinks.
7. Noisy or damaged electrodes that were previously removed were interpolated using spherical splines.
8. Data were segmented from -500 to 1,500ms relative to feedback stimuli.
9. Data were baseline corrected using a -200 to 0 ms time window.
10. Segments were removed if they violated 10 *µ*V/ms or 100 *µ*V max-min criteria.
11. Data were segmented to -200 to 1,000 ms relative to feedback stimuli.

#### Anti-Saccade

1. Noisy or faulty electrodes were identified using *EEGLAB’s clean rawdata* [1] algorithm using default parameters.
2. Data were filtered with a high pass filter using an FIR filter with a passband edge(s): 0.10Hz, filter order: 16500.00, cutoff frequency 0.05, and transition bandwidth: 0.10Hz.
3. *Zapline* [2] was applied to remove line noise artifacts, removing seven power line components.
4. Noisy or faulty channels were interpolated using spherical splines.
5. ICA was employed to identify and correct artifacts, including ocular artifacts, using *ICLabel* [1].
6. We then processed eye tracking data:
  - We identified saccades via an acceleration threshold of 8000° per second, a velocity threshold of 30° per second, and a deflection threshold of 0.1° (SR-Research [3]).
  - We identified fixations as intervals without saccades, which may include microsaccades.
  - We identified blinks when the pupil size was very small, and the pupil in the camera image was missing or severely distorted by eyelid occlusion.
7. EEG and eye tracking data were synchronized using *EYE EEG Toolbox* [4].
8. Data were reduced to 105 electrodes.
9. Data were re-referenced using an average reference.
10. Trials with incorrect performance (e.g., saccade moved in the wrong direction) and fast (*<* 100ms) or slow (*>* 800ms) saccade lengths were removed.
11. Data were baseline corrected using a -200 to 0ms time window.
12. data were segmented to -200 to 800 ms relative to the cue stimulus.

#### Face Perception & Visual Search

1. Data were downsampled to 256 Hz.
2. Data were re-referenced to an average reference.
3. DC offsets were removed.
4. Data were high-pass filtered at 0.1 Hz (order 2).
5. ICA was employed to identify and correct eye blinks and movements.
6. Data were segmented from -200 to 800 ms locked to image onset.
7. Data were baseline corrected using the -200 to 0 ms time period.
8. Excessively noisy channels were interpolated using spherical interpolation.
9. Segments were removed if they violated participant-specific artifact rejection thresholds.
10. Segments containing eye movements that survived ICA correction were identified and removed.
11. Segments with incorrect behavioural responses (e.g., incorrect performance, excessively fast reaction times) were removed.

### Benchmark Augmentation Techniques

#### Variational Autoencoder

A variational autoencoder (VAE) was used as a generative AI benchmark model as it is a leading generative method for augmenting EEG [5, 6, 7, 8, 9, 10, 11, 12, 13]. VAEs generate new samples by reducing high-dimensional data into probabilistic latent variables. They are composed of two networks: an Encoder *E*_*V AE*_ and a Decoder *D*_*V AE*_, each of which is a simple neural network.

We assume the input data *y* is generated from a probabilistic model according to the likelihood *p*(*y* | *z*) given the latent representation *z* ∼ *p*(*z*). Thus, the encoder *E*_*V AE*_ approximates the real posterior *p*(*z* | *y*) through variational inference with *q*(*z* | *y*; *θ*_*V AE*,*E*_) given the encoder parameters *θ*_*V AE*,*E*_. This way, the input data *y* is reduced to the sampled latent representation 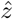. The decoder *D*_*V AE*_, on the other hand, approximates the likelihood 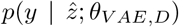 given the decoder parameters *θ*_*V AE*,*D*_ and thus approximates the input data *y* passed to the encoder with the reconstructed output *ŷ*.

Since the VAE is trained by performing variational inference, the objective can be expressed as the maximization of the evidence lower bound (ELBO) according to

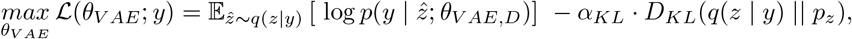

with the VAE model parameters *θ*_*V AE*_ = (*θ*_*V AE*,*E*_, *θ*_*V AE*,*D*_), the temperature coefficient *α*_*KL*_ and the Kullback-Leibler Divergence

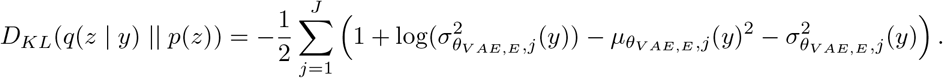

Hereby, the variables 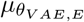 and 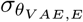 denote the mean and the variance parameters of the approximated posterior distribution *q*(*z* | *y*) which resembles a multivariate Gaussian distribution. The encoder generates these parameters to enable backpropagation of gradients according to the reparameterization trick

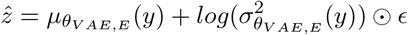

where *ϵ∼* 𝒩 (0, 1) and *⊙* signifying element-wise multiplication.

In line with the GAN, VAE training was implemented on trial-level EEG data and used an ADAM optimizer with a *l*_*r*_ = 0.0001 and decay rates of *β*_1_ = 0, *β*_2_ = 0.9.

### Transformation Augmentation Techniques

Unlike generative augmentation techniques, which produce additional samples to enhance classifier training, transformation augmentation techniques enhance classification performance by modifying empirical samples. Here, we have used five well-established EEG data augmentation techniques as benchmark methods: oversampling, Gaussian noise, smooth time masking, sign flip, and time reverse. Aside from oversampling, all of these were inspired by Rommel et al.’s [14] thorough investigations on standard augmentation techniques for enhancing EEG classification performance. From their research, we included all augmentation techniques that targeted the time-series dimension of EEG. These transformation augmentation techniques modify samples *y* in place, thus removing the original sample from the training dataset and replacing it with the transformed samples *ŷ*. An important hyper-parameter is the number of transformed samples *p*_*aug*_. In line with Rommel et al. [14], we set *p*_*aug*_ = 0.5 so that each sample had a 50% chance of being transformed.

#### Oversampling

Oversampling does not modify empirical samples directly but instead increases the strength of each sample by duplicating its existence within the training dataset. Our oversampling augmentations double the training datasets by duplicating each sample once. This method is akin to doubling the number of training epochs. The importance of this technique as a benchmark is to ensure that the GAN-augmentation effects are not simply a consequence of including additional training samples.

#### Gaussian Noise

Gaussian noise augmentation adds random noise to each data point within a sample. This technique increases classification performance as it makes the classifier more robust to noise. We applied a Gaussian noise with a *σ* = 0.1:

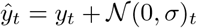

#### Smooth Time Masking

Smooth time masking removes a section of the sample by converting the values to zero. This technique increases the robustness of classifiers by forcing them to rely on the whole time-series data rather than specific segments that may be most predictive. To avoid sharp changes in the data between masked and non-masked segments, the masked segments were flanked by opposing sigmoid functions. Applying this technique requires a length of the mask *t*_Δ_ and the initial location where the mask begins *t*_*i*_. In this study, *t*_Δ_ was chosen from a uniform distribution ranging from 10% to 25% of the time-series length:

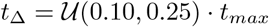

and *t*_*i*_ was chosen from a uniform distribution ranging from the first timepoint *t*_*min*_ to the last timepoint *t*_*max*_ minus the length of *t*_Δ_:

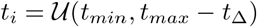

The smooth time mask is then here applied as:

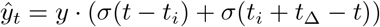

where *σ* is the sigmoid function:

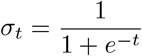

#### Time Reverse

Time reverse transformations flip the time domain to be backward. The flipping of the time domain conserves information about the sample in terms of the underlying frequencies and largely in terms of the signal waveforms. Thus, it enhances classification performance by providing unique samples that contain the same relative information. It is here applied as:

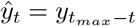

**Sign Flip** Sign flip augmentation reverses the sample’s polarity. This method increases a classifier’s robustness to noise and discriminative power as the transformed signals contain the same information but with a different representation. This method is applied as:

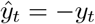

## Extended Tables

**Table S1.** Number of participants included for evaluation and classification train, validation, and test sets for each dataset.

|  | **Total** | **Evaluations** | **Train** | **Validation** | **Test** |
| --- | --- | --- | --- | --- | --- |
| **Reinforcement Learning** | 500 | 500 | 100 | 200 | 200 |
| **Anti-Saccade** | 199 | 189 | 100 | 49 | 50 |
| **Face Perception** | 40 | 37 | 20 | 10 | 10 |
| **Visual Search** | 40 | 35 | 20 | 10 | 10 |

**Table S2.** Hyperparameter configurations explored for each classifier: neural networks (NN), support vector machines (SVM), logistic regressions (LR), random forests (RF), and K-nearest neighbours (KNN) using Scikit-learn for evaluating augmentation impact on classification performance.

|  |  |  |
| --- | --- | --- |
| **NN** | Architectures | $N_1 = 25$<br>$N_1 = 50$<br>$N_1 = 25, N_2 = 25$<br>$N_1 = 50, N_2 = 50$<br>$N_1 = 50, N_2 = 25, N_3 = 50$ |
|  | L2 Regularization Strength | 0.05, 0.0001 |
|  | Activation Functions | Logistic, TanH, ReLU |
|  | Learning Rate Schedules | Constant, Inverse Scaling, Adaptive Learning |
|  | Maximum Iterations | 10,000; 20,000 |
| **SVM** | Inverse Regularization Strengths | 0.1, 1, 10, 100 |
|  | Kernel Types | Radial Basis, Polynomial, Sigmoid |
|  | Kernel Coefficients | 1.0, 0.1, 0.01, 0.001 |
| **LR** | Solver | liblinear |
|  | Penalties | L1, L2 |
| | Inverse Regularization Strengths | $10^{-4} \dots 10^4$, 20 steps |
|  | Maximum Iterations | 5,000; 10,000; 20,000; 50,000 |
| **RF** | Leaf Size | 5 |
|  | Number of Trees | 25, 50, 75, 100, 125, 150 |
|  | Maximum Depth | 3, 5, 7 |
| | Maximum Features | $\sqrt{N_{features}}$, $\log_2(N_{features})$ |
|  | Quality of Split Criterion | Gini Impurity, Entropy |
| **KNN** | Number of Neighbors | $SS = \begin{cases} 1-10 & \Rightarrow 3, 5, 7 \\ 11-20 & \Rightarrow 6, 10, 14 \\ 21-30 & \Rightarrow 9, 15, 21 \\ \geq 31 & \Rightarrow 12, 18, 28 \end{cases}$ |
|  | Weight Function | Uniform, Distance |
|  | Distance Metric | Euclidean, Manhattan, Minkowski |

## Extended Figures

**Fig. S1.**
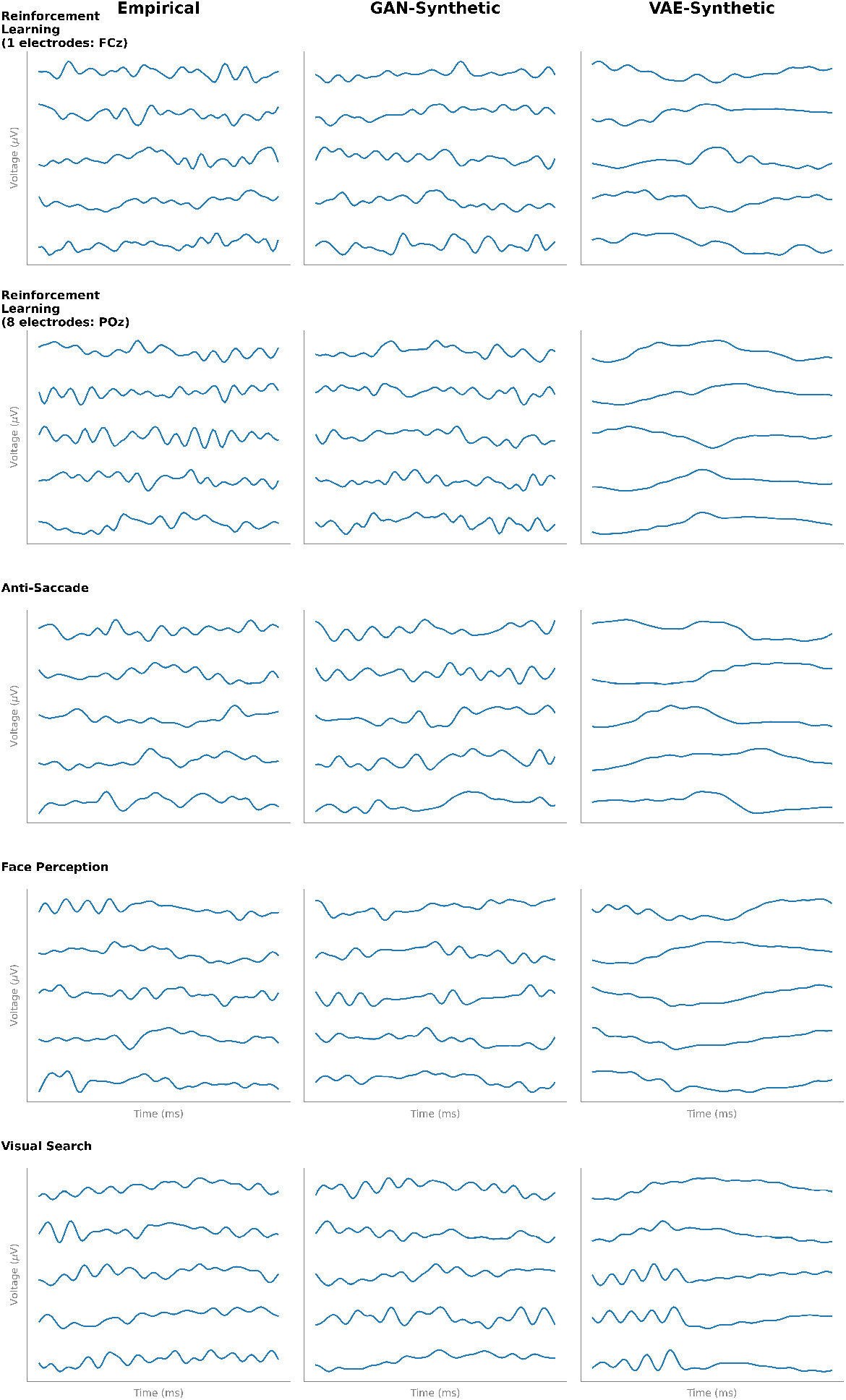
Samples of empirical and synthetically generated single-trial EEG data across four datasets.

**Fig. S2.**
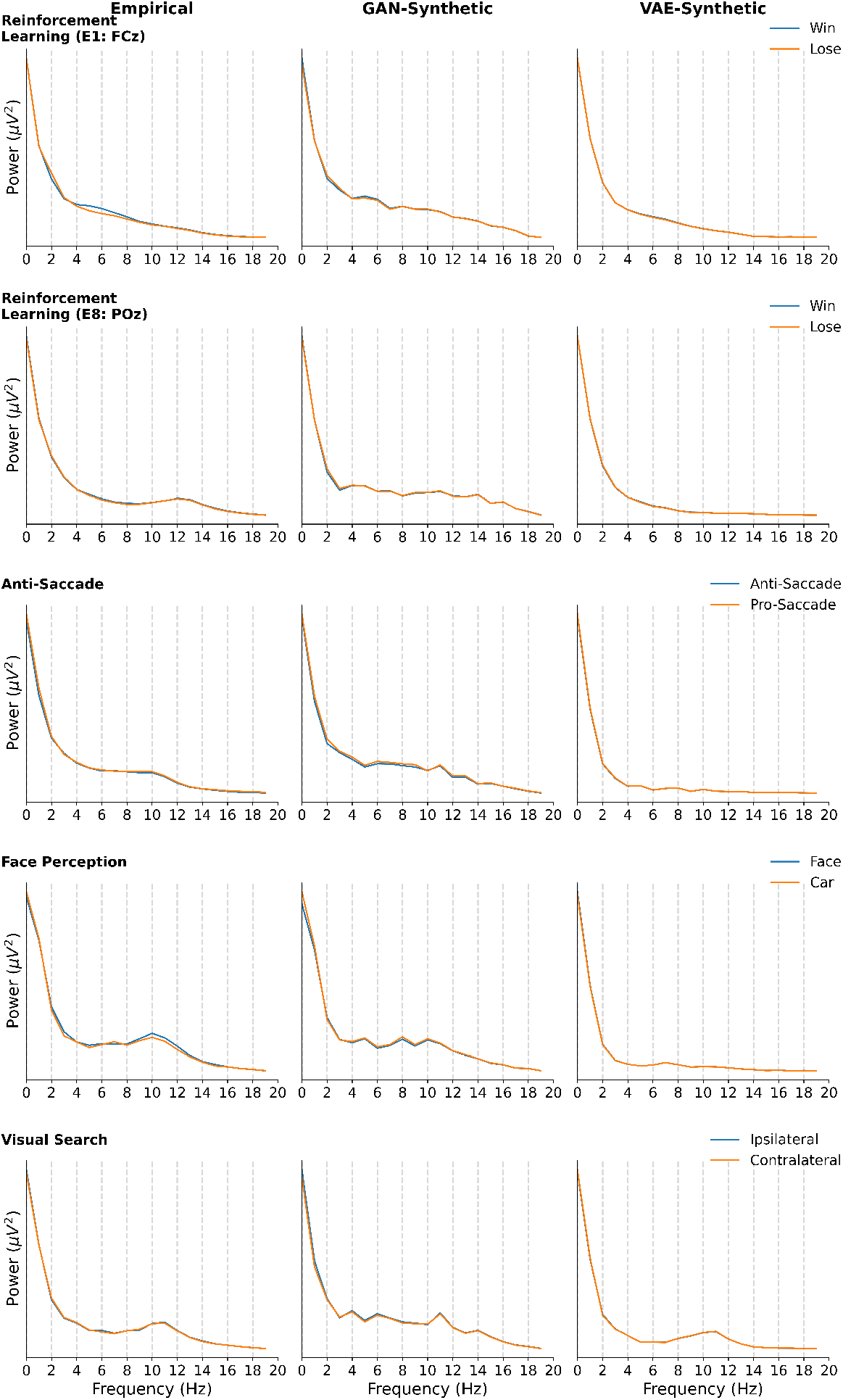
Grand-averaged conditional frequency (FFT) data from empirical and synthetic samples produced by the GAN and VAE models.

**Fig. S3.**
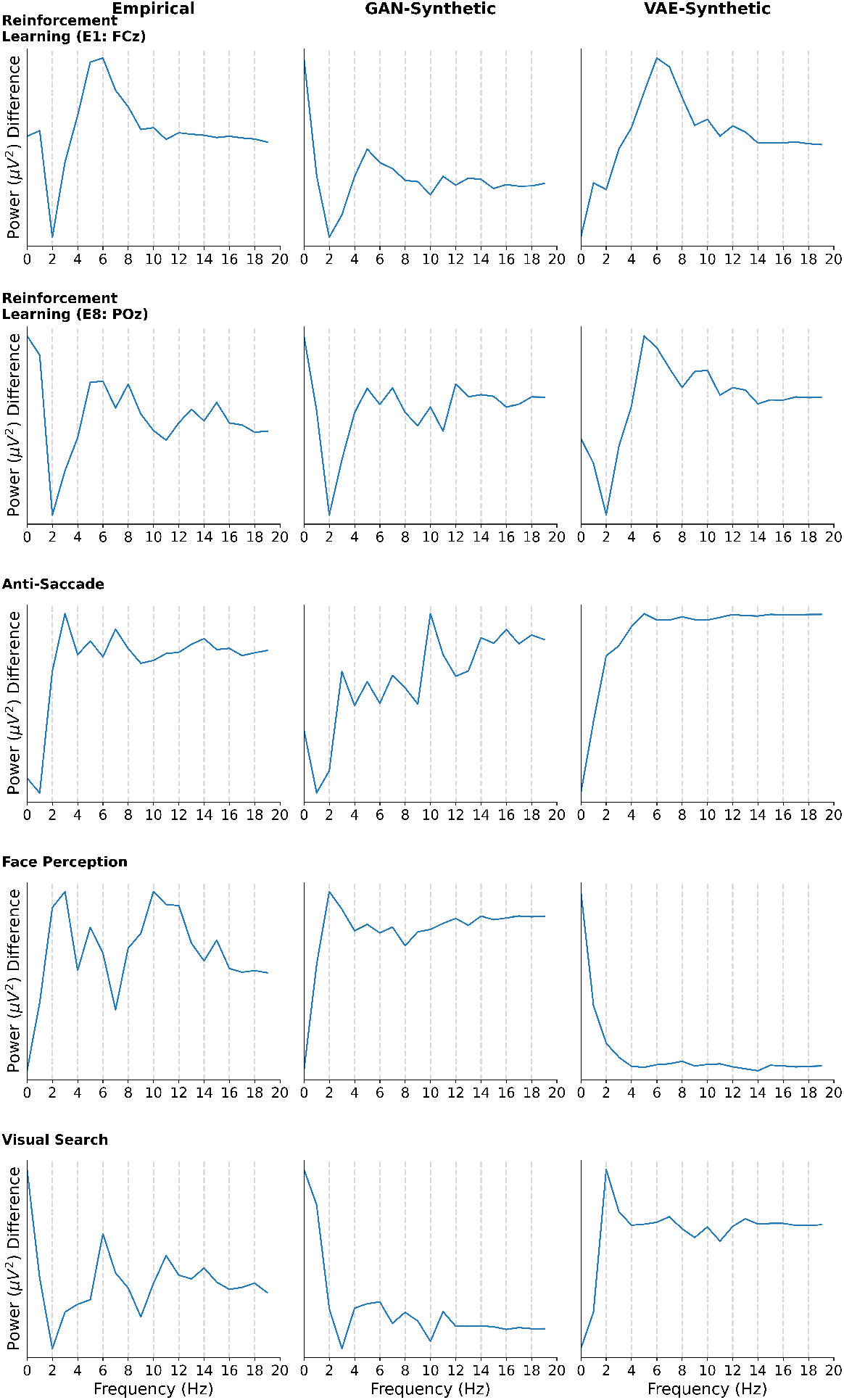
Grand-averaged difference frequency (FFT) data from empirical and synthetic samples produced by the GAN and VAE models.

**Fig. S4.**
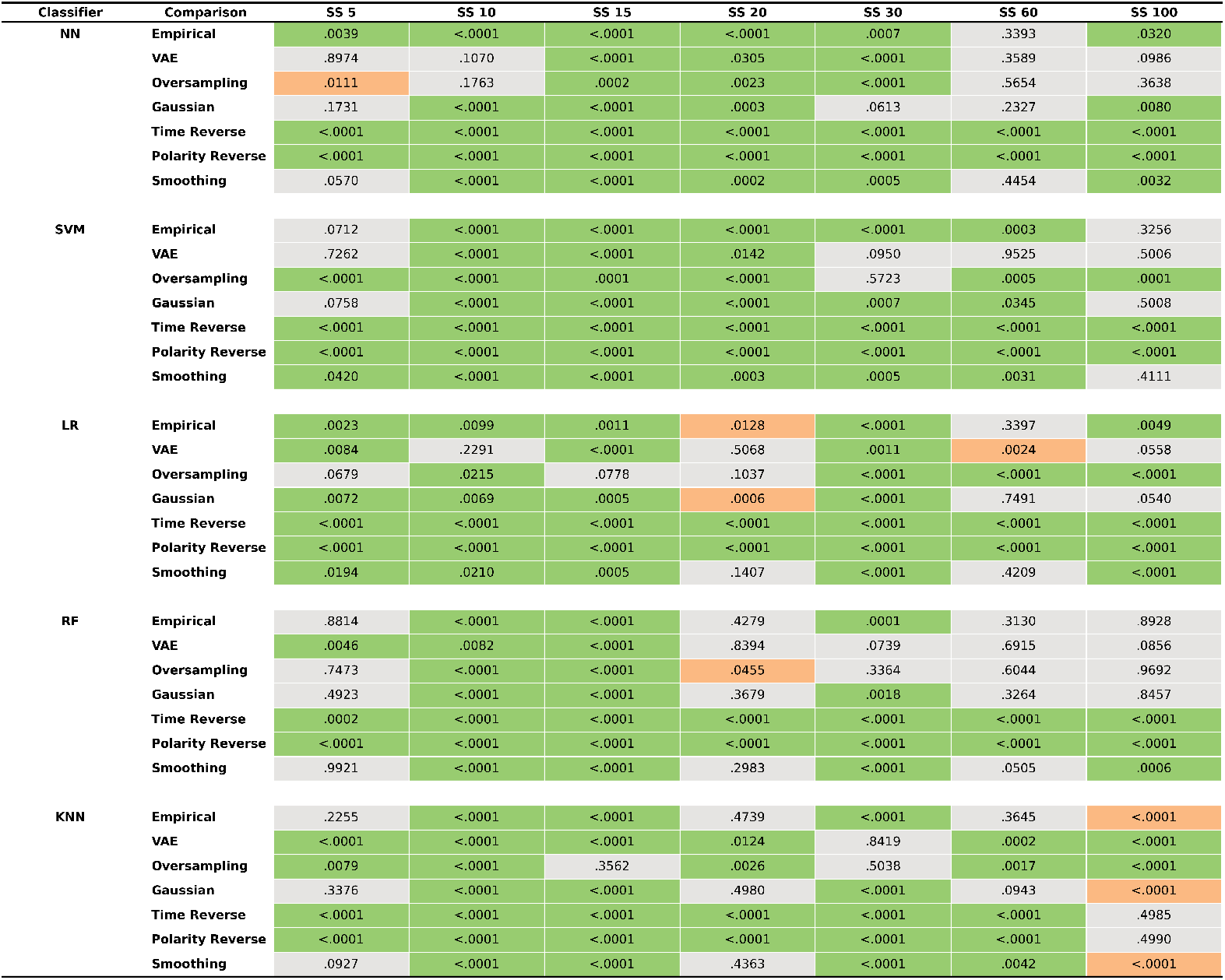
Bootstrap results of the reinforcement learning (1 electrode) dataset for the comparison of GAN performance with empirical and benchmark performance across classifiers and sample sizes. Values presented are the p-values of the mean difference. Green values indicate a significant increase in performance for the GAN relative to the corresponding comparison dataset, gray values indicate a non-significant difference in performance between the GAN and the comparison dataset, and red values indicate a significant decrease in performance for the GAN relative to the corresponding comparison dataset. SS = Sample Size, NN = Neural Network, SVM = Support Vector Machine, LR = Logistic Regression, RF = Random Forest, KNN = K-Nearest Neighbors.

**Fig. S5.**
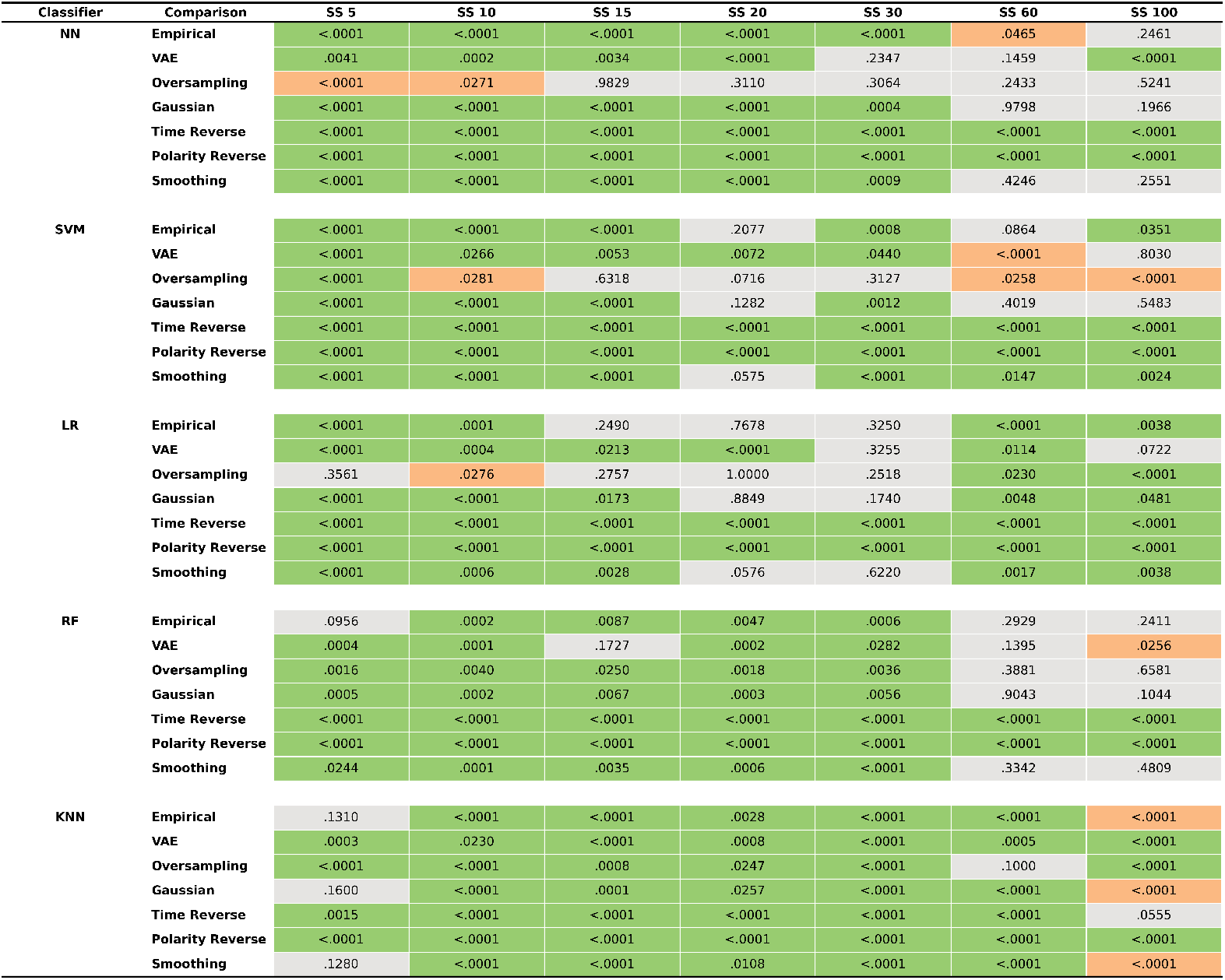
Bootstrap results of the reinforcement learning (8 electrode) dataset for the comparison of GAN performance with empirical and benchmark performance across classifiers and sample sizes. Values presented are the p-values of the mean difference. Green values indicate a significant increase in performance for the GAN relative to the corresponding comparison dataset, gray values indicate a non-significant difference in performance between the GAN and the comparison dataset, and red values indicate a significant decrease in performance for the GAN relative to the corresponding comparison dataset. SS = Sample Size, NN = Neural Network, SVM = Support Vector Machine, LR = Logistic Regression, RF = Random Forest, KNN = K-Nearest Neighbors.

**Fig. S6.**
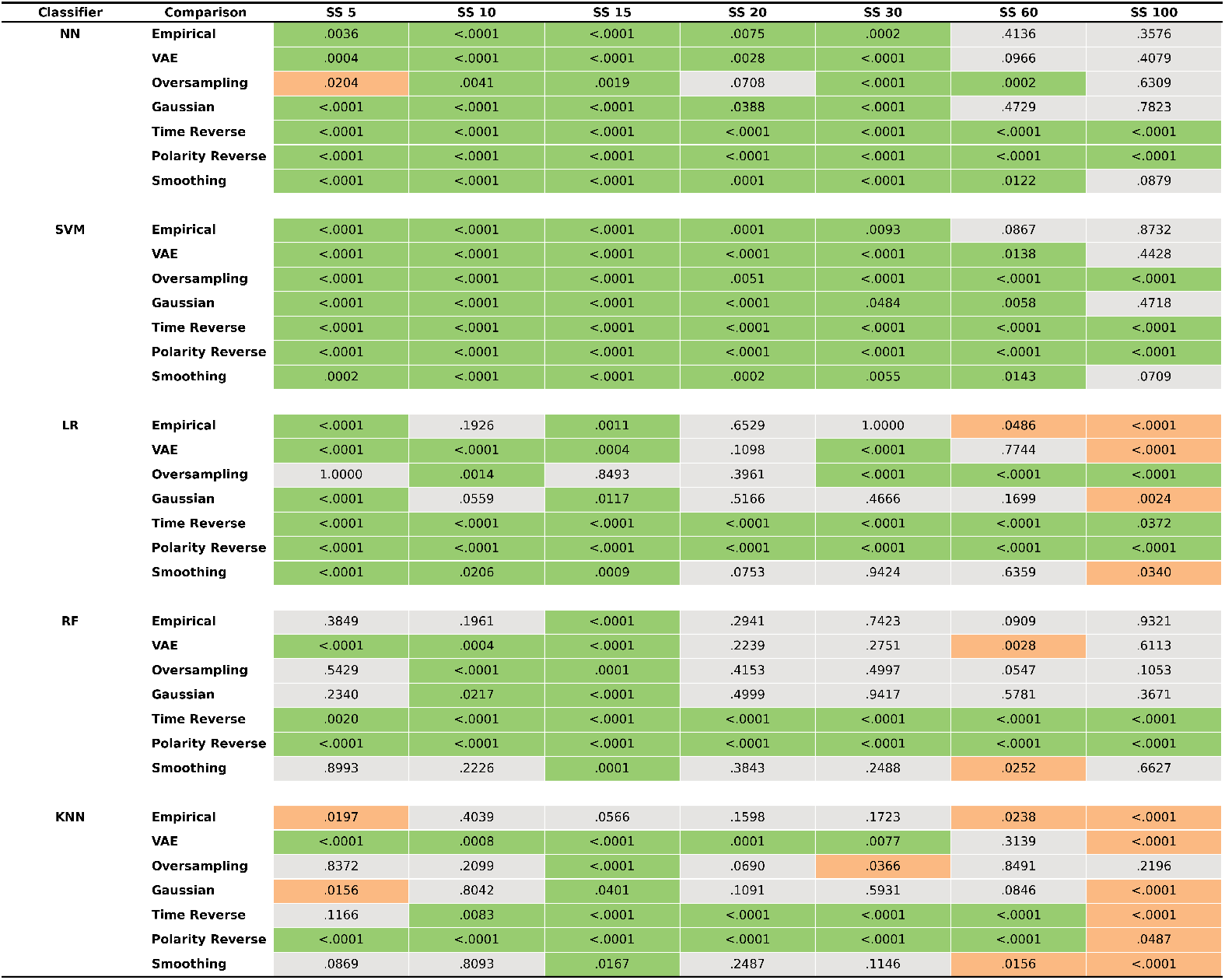
Bootstrap results of the anti-saccade dataset for the comparison of GAN performance with empirical and benchmark performance across classifiers and sample sizes. Values presented are the p-values of the mean difference. Green values indicate a significant increase in performance for the GAN relative to the corresponding comparison dataset, gray values indicate a non-significant difference in performance between the GAN and the comparison dataset, and red values indicate a significant decrease in performance for the GAN relative to the corresponding comparison dataset. SS = Sample Size, NN = Neural Network, SVM = Support Vector Machine, LR = Logistic Regression, RF = Random Forest, KNN = K-Nearest Neighbors.

**Fig. S7.**
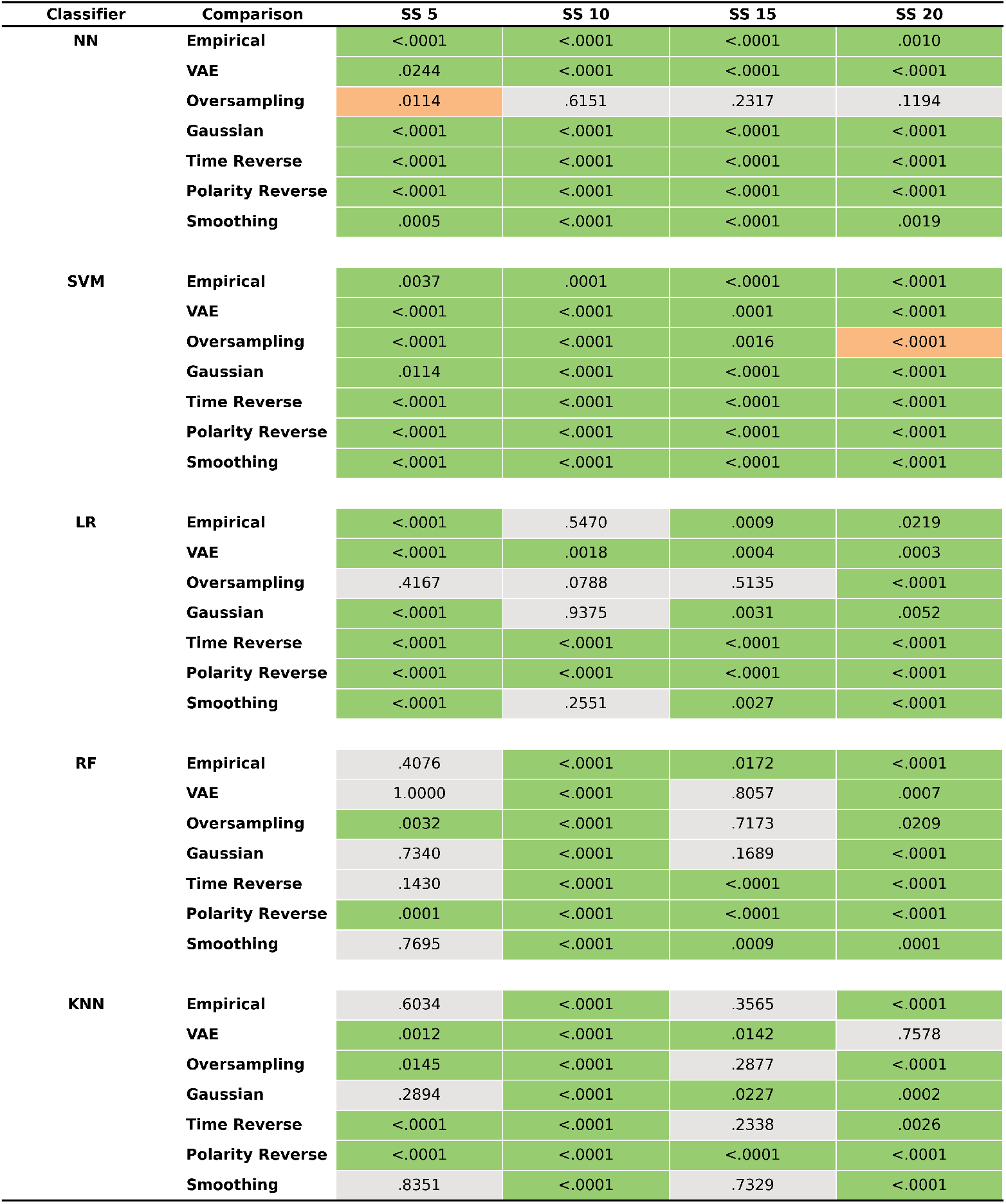
Bootstrap results of the face processing dataset for the comparison of GAN performance with empirical and benchmark performance across classifiers and sample sizes. Values presented are the p-values of the mean difference. Green values indicate a significant increase in performance for the GAN relative to the corresponding comparison dataset, gray values indicate a non-significant difference in performance between the GAN and the comparison dataset, and red values indicate a significant decrease in performance for the GAN relative to the corresponding comparison dataset. SS = Sample Size, NN = Neural Network, SVM = Support Vector Machine, LR = Logistic Regression, RF = Random Forest, KNN = K-Nearest Neighbors.

**Fig. S8.**
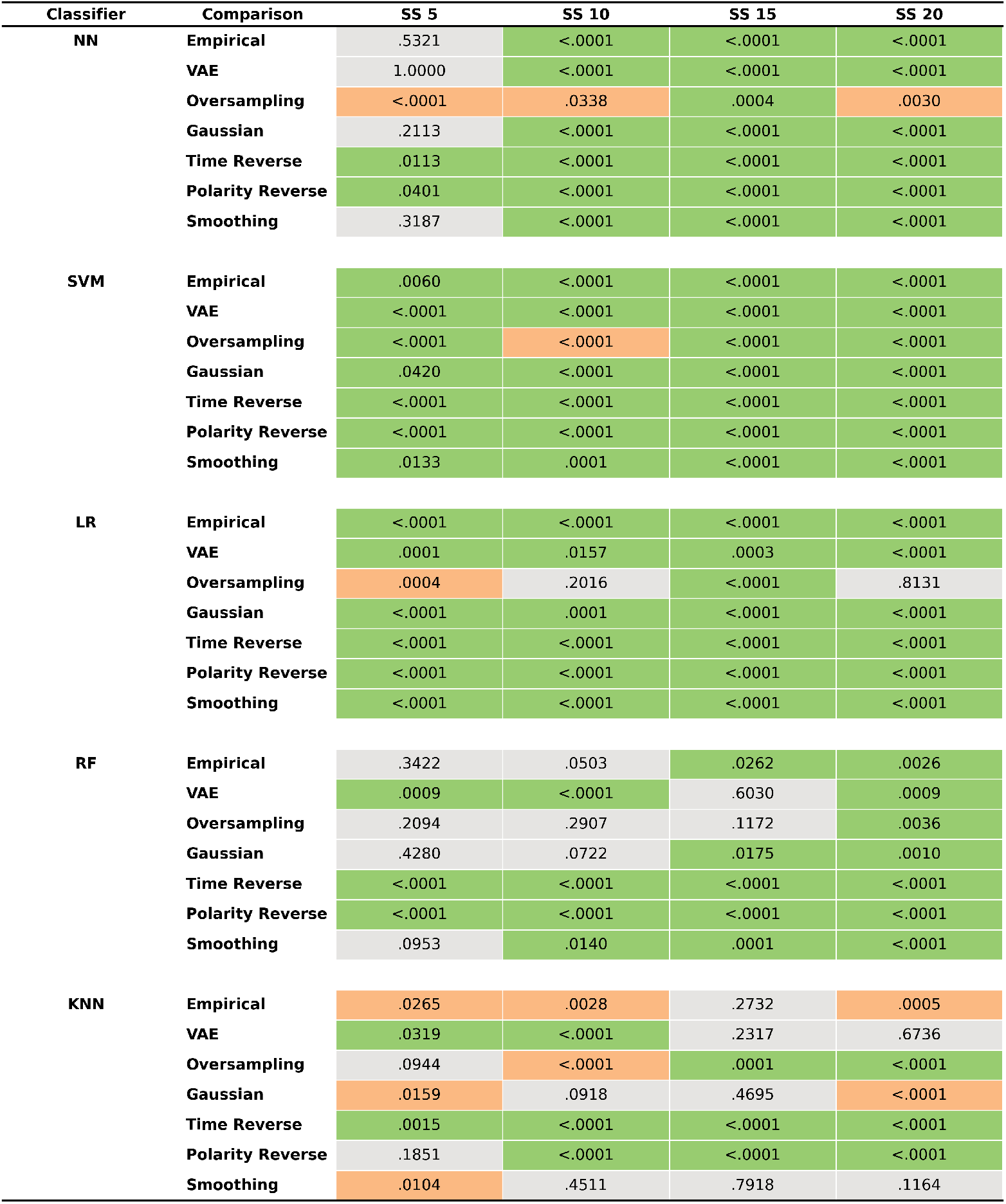
Bootstrap results of the visual search dataset for the comparison of GAN performance with empirical and benchmark performance across classifiers and sample sizes. Values presented are the p-values of the mean difference. Green values indicate a significant increase in performance for the GAN relative to the corresponding comparison dataset, gray values indicate a non-significant difference in performance between the GAN and the comparison dataset, and red values indicate a significant decrease in performance for the GAN relative to the corresponding comparison dataset. SS = Sample Size, NN = Neural Network, SVM = Support Vector Machine, LR = Logistic Regression, RF = Random Forest, KNN = K-Nearest Neighbors.

**Fig. S9.**
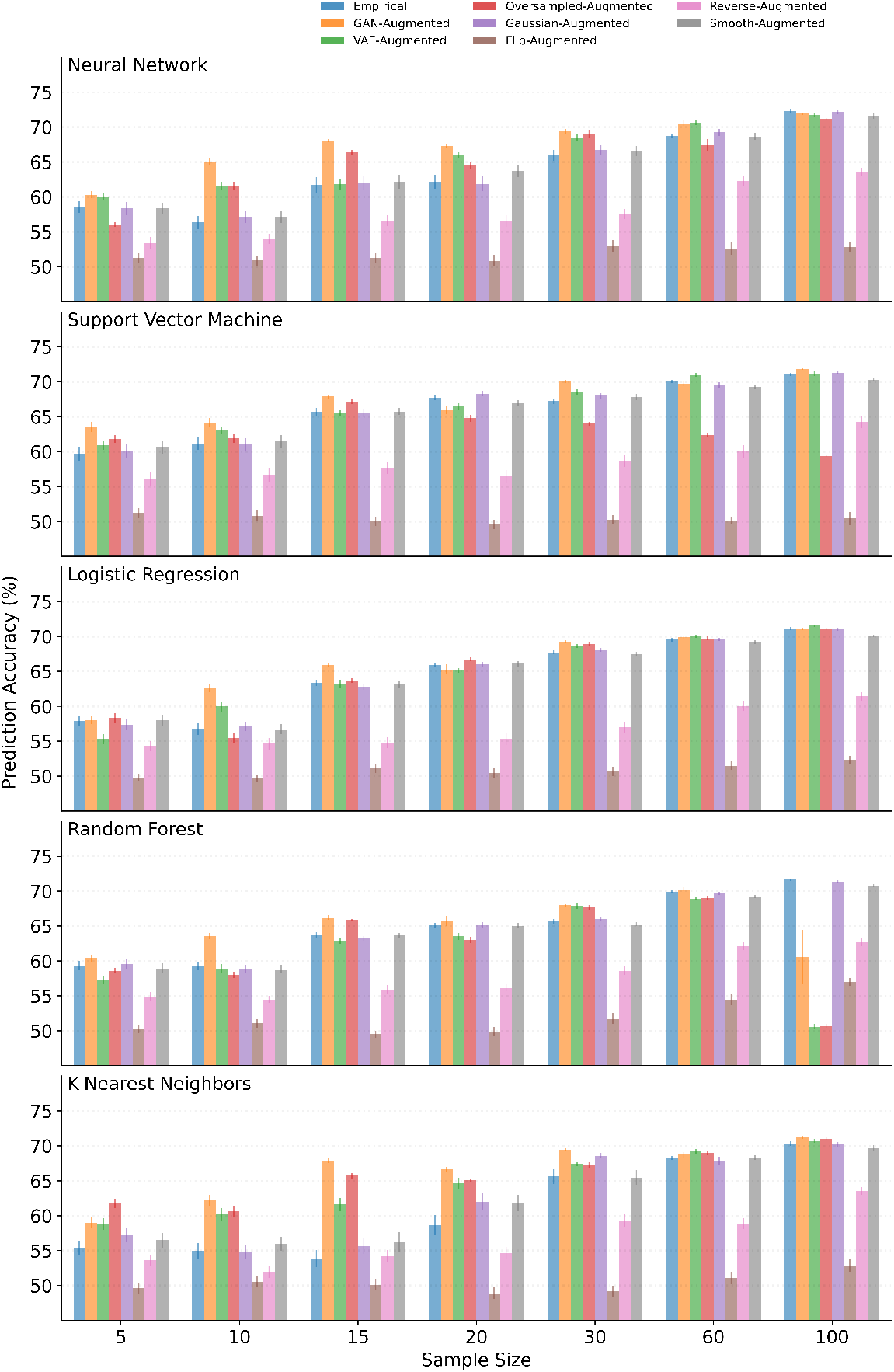
2-way classification performance (%) achieved via GAN augmentation with six benchmark techniques and empirical data across sample sizes for the reinforcement learning dataset containing one electrode. Findings suggest that the GAN robustly outperforms other methods in enhancing EEG sample quality across all classifiers and sample sizes. Error bars represent standard error of the mean across participants.

**Fig. S10.**
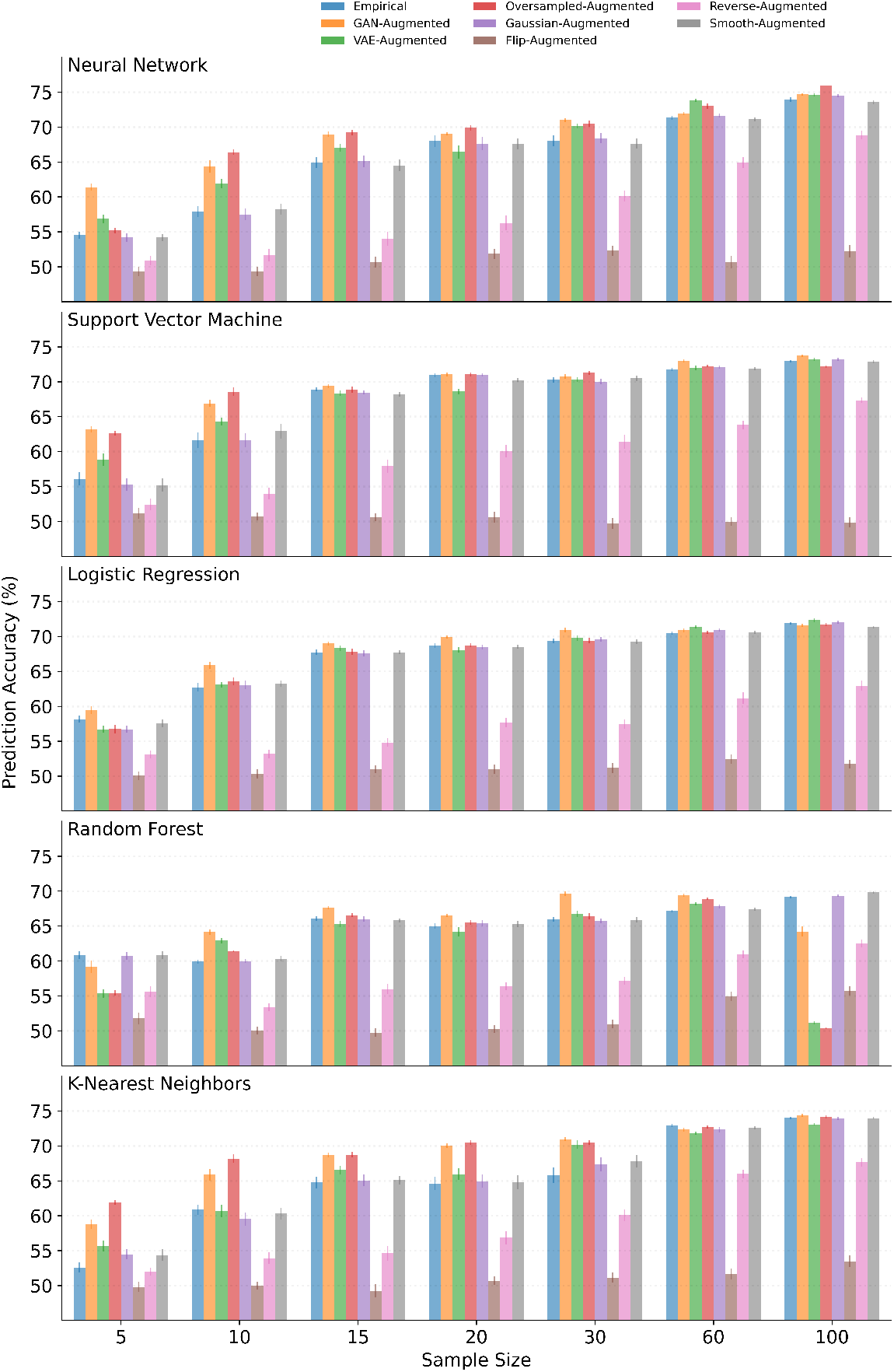
2-way classification performance (%) of GAN augmentation compared with six benchmark techniques and empirical data across sample sizes for the reinforcement learning dataset containing eight electrodes. Findings suggest that the GAN robustly outperforms other methods in enhancing EEG sample quality across all classifiers and sample sizes. Error bars represent standard error of the mean across participants.

**Fig. S11.**
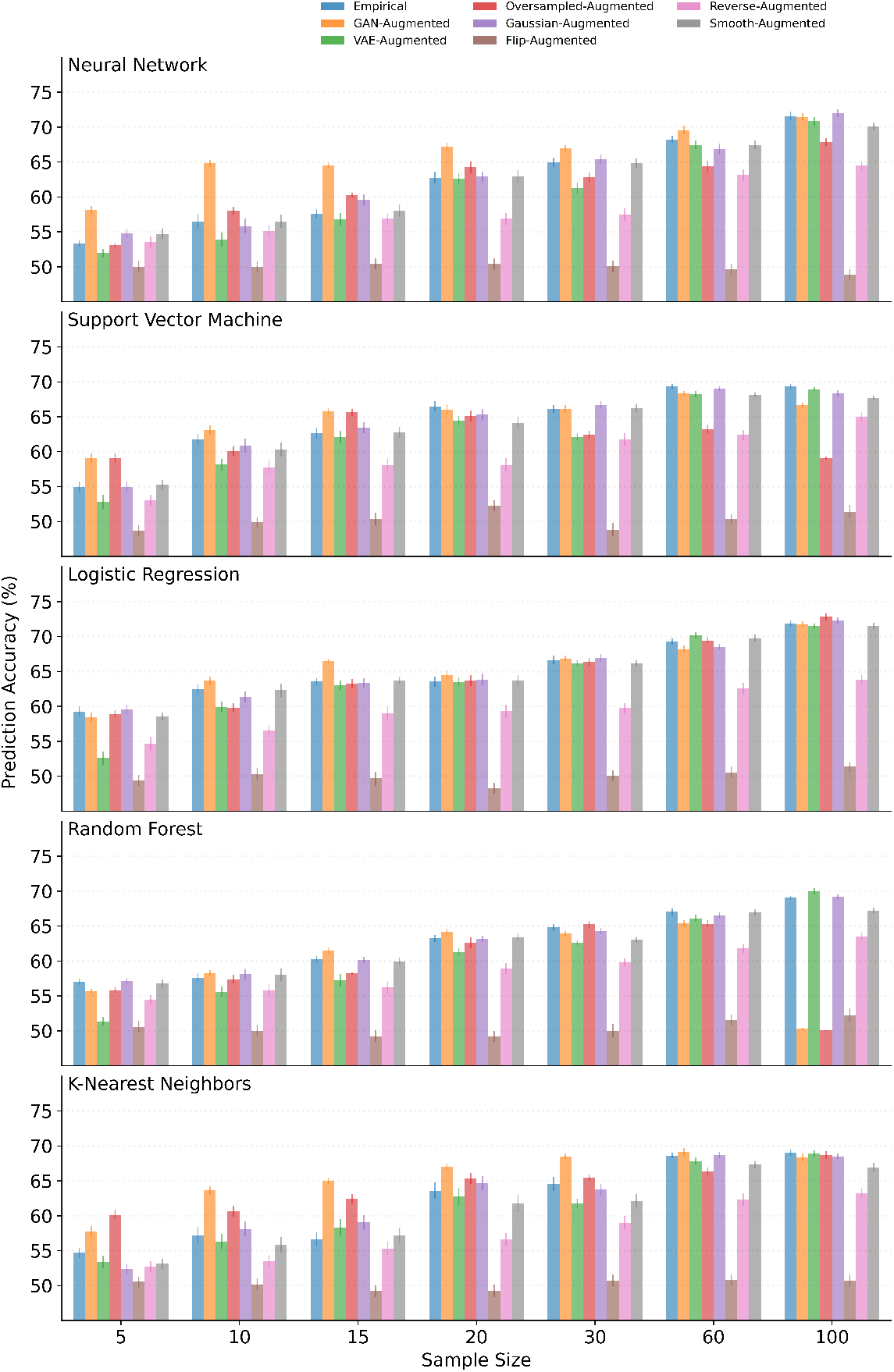
2-way classification performance (%) of GAN augmentation compared with six benchmark techniques and empirical data across sample sizes for the face anti-saccade dataset. Findings suggest that the GAN robustly outperforms other methods in enhancing EEG sample quality across all classifiers and sample sizes. Error bars represent standard error of the mean across participants.

**Fig. S12.**
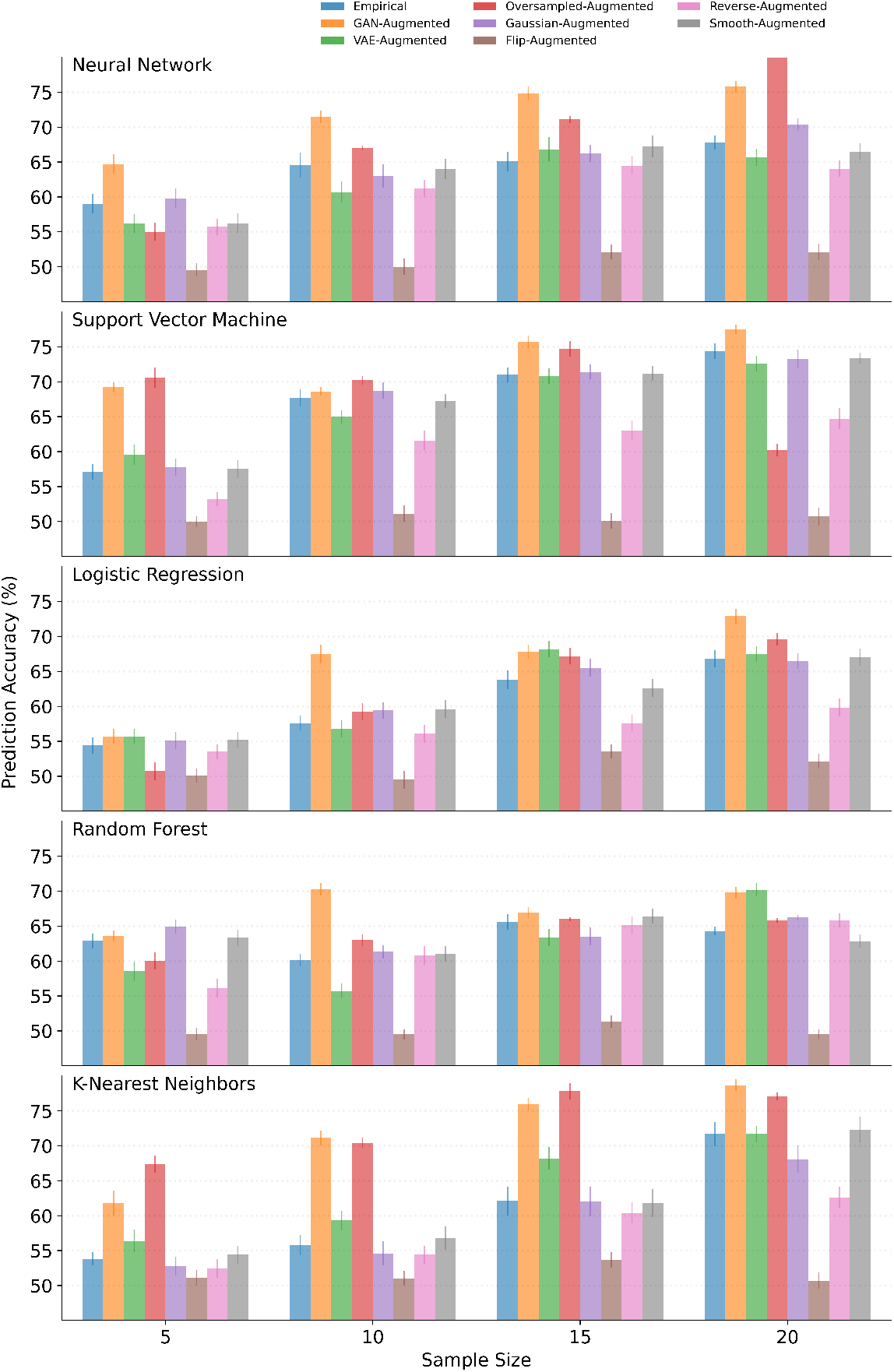
2-way classification performance (%) of GAN augmentation with six benchmark techniques and empirical data across sample sizes for the face perception dataset. Findings suggest that the GAN robustly outperforms other methods in enhancing EEG sample quality across all classifiers and sample sizes. Error bars represent standard error of the mean across participants.

**Fig. S13.**
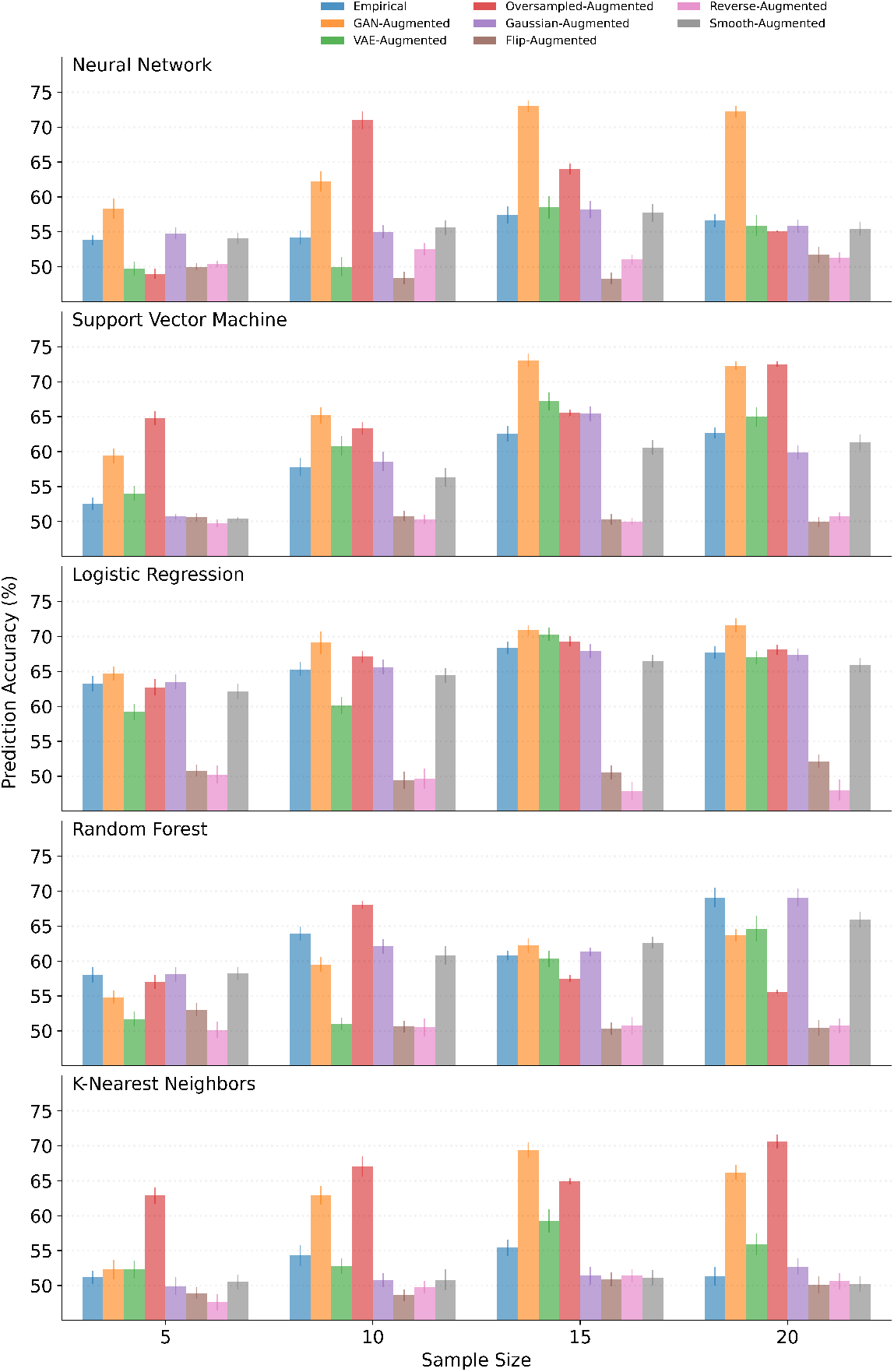
2-way classification performance (%) of GAN augmentation with six benchmark techniques and empirical data across sample sizes for the visual search dataset. Findings suggest that the GAN robustly outperforms other methods in enhancing EEG sample quality across all classifiers and sample sizes. Error bars represent standard error of the mean across participants.

